# The human ciliopathy protein RSG1 links the CPLANE complex to transition zone architecture

**DOI:** 10.1101/2024.09.25.614984

**Authors:** Neftalí Vazquez, Chanjae Lee, Irene Valenzuela, Thao P. Phan, Camille Derderian, Marcelo Chávez, Nancie A. Mooney, Janos Demeter, Mohammad Ovais Aziz-Zanjani, Ivon Cusco, Marta Codina, Núria Martínez-Gil, Diana Valverde, Carlos Solarat, Ange-Line Buel, Cristel Thauvin-Robinet, Elisabeth Steichen, Isabel Filges, Pascal Joset, Julie De Geyter, Krishna Vaidyanathan, Tynan Gardner, Michinori Toriyama, Edward M. Marcotte, Elle C. Roberson, Peter K. Jackson, Jeremy F. Reiter, Eduardo F. Tizzano, John B. Wallingford

## Abstract

Cilia are essential organelles and variants in genes governing ciliary function result in ciliopathic diseases. The Ciliogenesis and PLANar polarity Effectors (CPLANE) protein complex is essential for ciliogenesis in animals models but remains poorly defined. Notably, all but one subunit of the CPLANE complex have been implicated in human ciliopathy. Here, we identify three families in which variants in the remaining CPLANE subunit *CPLANE2/RSG1* also cause ciliopathy. These patients display cleft palate, tongue lobulations and polydactyly, phenotypes characteristic of Oral-Facial-Digital Syndrome. We further show that these alleles disrupt two vital steps of ciliogenesis, basal body docking and recruitment of intraflagellar transport proteins. Moreover, APMS reveals that Rsg1 binds the CPLANE and also the transition zone protein Fam92 in a GTP-dependent manner. Finally, we show that CPLANE is generally required for normal transition zone architecture. Our work demonstrates that CPLANE2/RSG1 is a causative gene for human ciliopathy and also sheds new light on the mechanisms of ciliary transition zone assembly.

## Introduction

Ciliopathies are a broad class of human diseases that share an etiology of defective cilia structure or function. These diseases span skeletal anomalies, craniofacial defects, cystic kidneys, blindness, obesity and other clinical manifestations, highlighting the wide array of physiological functions that require components of the cilium ^1–3^. Cilia are assembled and maintained by a cohort of multiprotein machines, such as the IFT complexes ^4,5^, the BBSome ^6^, transition zone complexes ^7^, and variants in subunits of these complexes are sufficient to cause ciliopathy ^2,3^. The Ciliogenesis and PLANar polarity Effector (CPLANE) complex is also essential for ciliogenesis in all vertebrates including humans ^8^, yet its function remains far less well defined.

Identified initially as a tripartite complex that controls planar cell polarity in *Drosophila*, the vertebrate CPLANE complex comprises five proteins ^8^. Fuz/Cplane3 and Intu/Cplane4 were the first vertebrate orthologues to be described; in *Xenopus* they are essential for ciliogenesis by dint of their roles in basal body docking and recruitment of IFT-A2 proteins to the base of cilia ^9–12^. More recently, it was shown that Fuz and Intu together form a GEF for Rab23, which in turn is implicated in the docking of basal bodies to the apical surface _13._

Rsg1/Cplane2 was identified as a Fuz-interacting small GTPase essential for ciliogenesis in *Xenopus* ^12,14^. Recent work suggests it is not a substrate of the Intu/Fuz GEF, but rather an effector ^15^. Wdpcp/Cplane5 encodes a beta-propeller protein and was first found to be essential for ciliogenesis in *Xenopus* ^16^. Further studies revealed that all of these CPLANE subunits are essential for ciliogenesis in mice ^14,17–20^, with Rsg1 acting at a relatively late step in ciliogenesis ^21^.

Proteomic analysis revealed that Intu, Fuz, Wdpcp, and Rsg1 form a stable and discrete complex that also contains the human ciliopathy protein Jbts17/CPLANE1 ^22^. Recently, the structure of a partial CPLANE complex lacking Jbts17 was solved by CryoEM (**Fig. 1A**) revealing a similar structure to other hexa-longin domain GEFs such as Mon1-Czz1 and Hps1-Hps4 and that Fuz interacts directly with Rsg1^15^. Studies in a variety of cell types indicate that the CPLANE complex localizes near basal bodies, where it assembles hierarchically, with Rsg1 being the most downstream component ^21,22^.

**Fig 1.**
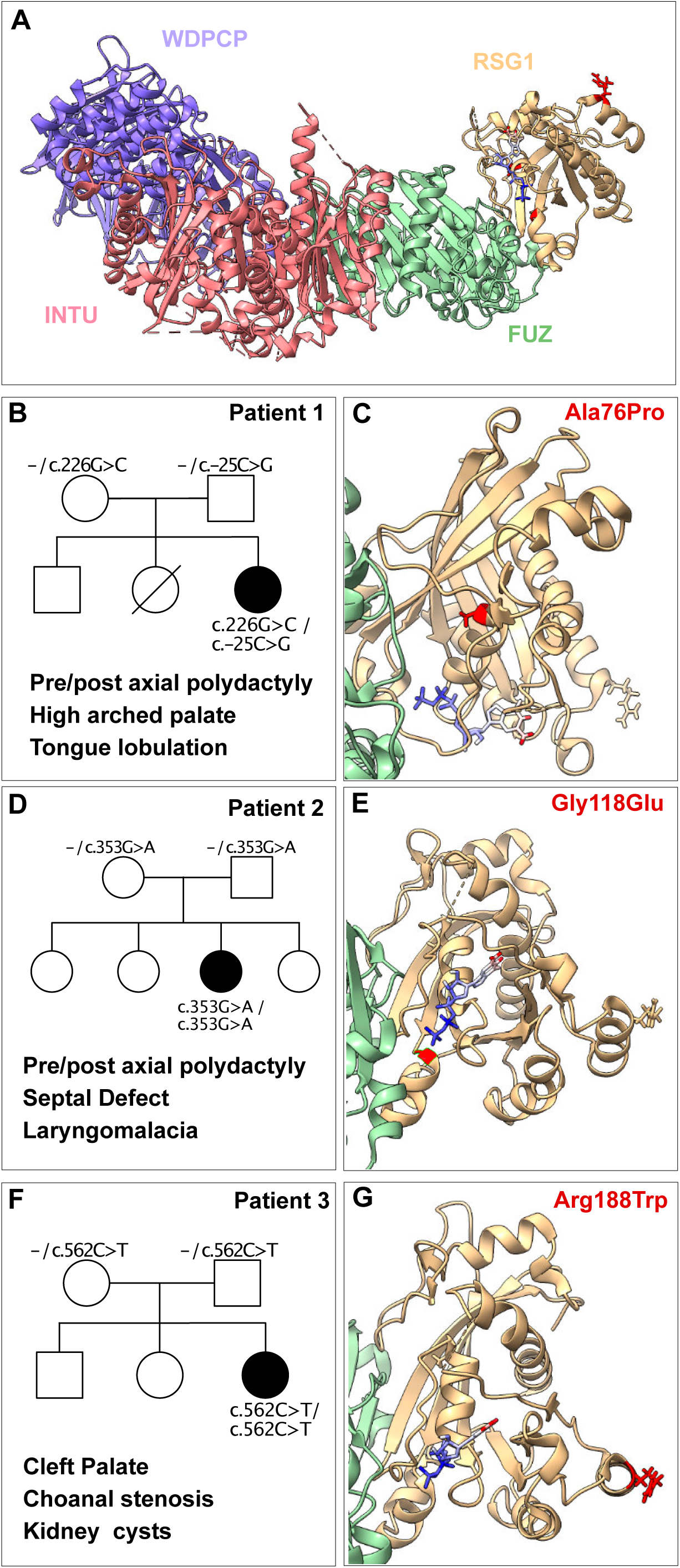
Allelic variants of *RSG1/CPLANE2* are associated with human ciliopathy. (A) Structure of mouse CPLANE complex, Wdpcp, Intu, Fuz, and Rsg1 (PDB: 7Q3E). *RSG1* allelic variants highlighted in red and GTP in the nucleotide pocket indicated (B,D,F) Pedigree maps and prominent features for ciliopathy indicated patients. (C,E,G) Structure of mouse Rsg1 (PDB 7q3e) with corresponding human residues altered by ciliopathy variants highlighted in red (see also Supp. Fig. 2B, C).

Finally, pathogenic variants in all but one CPLANE subunit have now been shown to be causative for human ciliopathy ^16,22–29^. These variants disrupt both lipid binding by the CPLANE complex and IFT-A2 recruitment to basal bodies ^15,22^. Though disruption of basal body docking is a common feature of CPLANE disruption in *Xenopus* ^10,12^, it remains unknown if ciliopathic variants in human *CPLANE* genes affect basal body docking. Unlike all other CPLANE genes, *CPLANE2* has yet to be linked to human disease.

Here, we identified three families in which homozygous or compound heterozygous variants in *CPLANE2* correlate with a ciliopathy phenotype within the spectrum of Oral-Facial-Digital syndrome (OFD). More precisely, the phenotype was quite similar to that reported for another of the ciliopathies related to the CPLANE complex (*CPLANE1* ^23^). Using *in vivo* imaging, we show that these alleles disrupt not only IFT-A2 recruitment but also basal body docking. One of these alleles lies within the GTP-binding domain of Rsg1, and proteomic analysis revealed that Rsg1 interaction with all CPLANE subunits is GTP-dependent. We also discovered a GTP-dependent interaction of Rsg1 and the BAR domain ciliopathy protein Fam92a. Based on that insight, we identified an unexpected role for Rsg1 and CPLANE in maintaining the normal architecture of the ciliary transition zone. Together, these results shed new light on the mechanisms of CPLANE action during ciliogenesis and how it is disrupted in CPLANE-associated ciliopathy.

## Results and Discussion

### Allelic variants of *CPLANE2/RSG1* cause human ciliopathy

We identified two patients in which variants in *CPLANE2* segregated with a spectrum of anomalies resembling OFD. The first patient presented with polyhydramnios during pregnancy, bilateral pre-and post-axial polydactyly on hands and feet, hypertelorism, high arched palate, and tongue lobulation (**Fig. 1B; Supplementary Clinical Report; Supp. Table 1**). This patient was previously described, but no molecular explanation was identified at that time ^30^. Exome sequencing identified compound heterozygous variants in *CPLANE2*. The maternally inherited variant (NM_030907.4:c.226G>C/c.-25C>G) lies in the coding region and changes alanine to proline at position 76, the impact of which is described below. The paternal allele (NM_030907.4:c.-25C>G) lies in the 5’ untranslated region. Analysis using RiboNN ^31^ predicts this allele will cause a modest but significant reduction in translation efficiency and thus affect RSG1 protein levels. (Note: To accommodate changing nomenclature, this paper will use the current formal human gene names (*CPLANE2*, etc.), but will use the names of protein subunits that are consistent with previously published literature (Rsg1, etc.).)

The second patient had prenatal detection of aortic coarctation and presented with a normal palate but a lobulated tongue and laryngomalacia, as well as a cardiac septal defect and post-axial polydactyly in one hand and pre-axial polydactyly on both feet. Exome sequencing for this patient identified a homozygous variant (NM_030907:c.G353A) in the coding region changing Gly 118 to Glu (**Fig. D; Supplementary Clinical Report; Supp. Table 1**).

Finally in the third patient, we observed a somewhat milder phenotype but also consistent with the clinical manifestations of the ciliopathy spectrum. During pregnancy, ultrasound identified hypoplastic and cystic dysplastic kidneys, oligohydramnios, microcephaly and IUGR. After birth the patient displayed no polydactyly, but presented with a cleft palate, choanal stenosis, pulmonary hypoplasia, and kidney cysts. In this patient, exome sequencing identified a homozygous variant (NM_030907:c.562C>T) that results in an Arg188Trp change in the coding region (**Fig. 1F; Supp. Fig. 1; Supplementary Clinical Report; Supp. Table 1**).

Because polydactyly and defects in the palate, tongue, larynx, and kidneys are hallmarks of CPLANE disruption in mice ^14,17–19,21,22,32^, these data raise the possibility that these variants in *CPLANE2/RSG1* are causative for the ciliopathy phenotype in these patients.

### Ciliopathy alleles disrupt RSG1 function via distinct mechanisms

To understand the molecular etiology of ciliopathic *CPLANE2* alleles, we first mapped their position to the known structure of RSG1 ^15^, which is generally similar to that of Rab GTPases (**Supp. Fig. 2**). AlphaFold3 ^33^ predicts human RSG1 to fold in a manner very similar to the known structure of mouse Rsg1, characterized by a central β-sheet surrounded by five α-helices (**Supp. Fig. 2B, C**). Mapping variant residues onto the predicted structure revealed that and A76 anchors the α1 helix; substitution of proline would be expected to severely disrupt the formation of that helix (**Fig. 1C; Supp. Fig. 2A, C**). G118 lies within the G3 region of the GTP binding pocket, immediately adjacent to the a key residue predicted to mediate GTP binding of Rsg1, E119^15^ (**Fig. 1E; Supp. Fig 1A, C**). By contrast, R188 lies on the outside edge of the α4 helix, away from the GTP-binding region and away from the known sites of Rsg1 interaction with Fuz (**Fig. 1G, Supp. Fig. 2A, C**).

To further explore pathogenicity, we tested these alleles’ effect on localization of Rsg1 protein using multiciliated cells (MCCs) in *Xenopus* (**Fig. 2A**), a proven platform for modeling the cell biology of ciliopathies ^34,35^. Because protein/protein interactions co-evolve, we used equivalent variants in the *Xenopus* orthologue of Rsg1 for these experiments (**Supp. Fig. 3**). As expected^14^, wild-type Rsg1 localized strongly to basal bodies (BB’s)(**Fig. 2B**), while in contrast, S72P (= human A76P) and G114E (= G118E) displayed significantly reduced enrichment at basal bodies (**Fig. 2C, D**). The S72P allele also appeared poorly expressed (**Fig. 2C**), and western blots confirmed that this variant severely destabilized the Rsg1 protein (**Supp. Fig. 4**). By contrast, while lost from basal bodies, the G114E allele appeared slightly enriched in the cytoplasm (not shown), and western blots indicated normal amounts of protein (**Supp. Fig. 4**). We quantified these alleles’ reduced recruitment to BBs as the ratio of Rsg1-GFP to Centrin-RFP (**Fig. 2F**).

**Fig 2.**
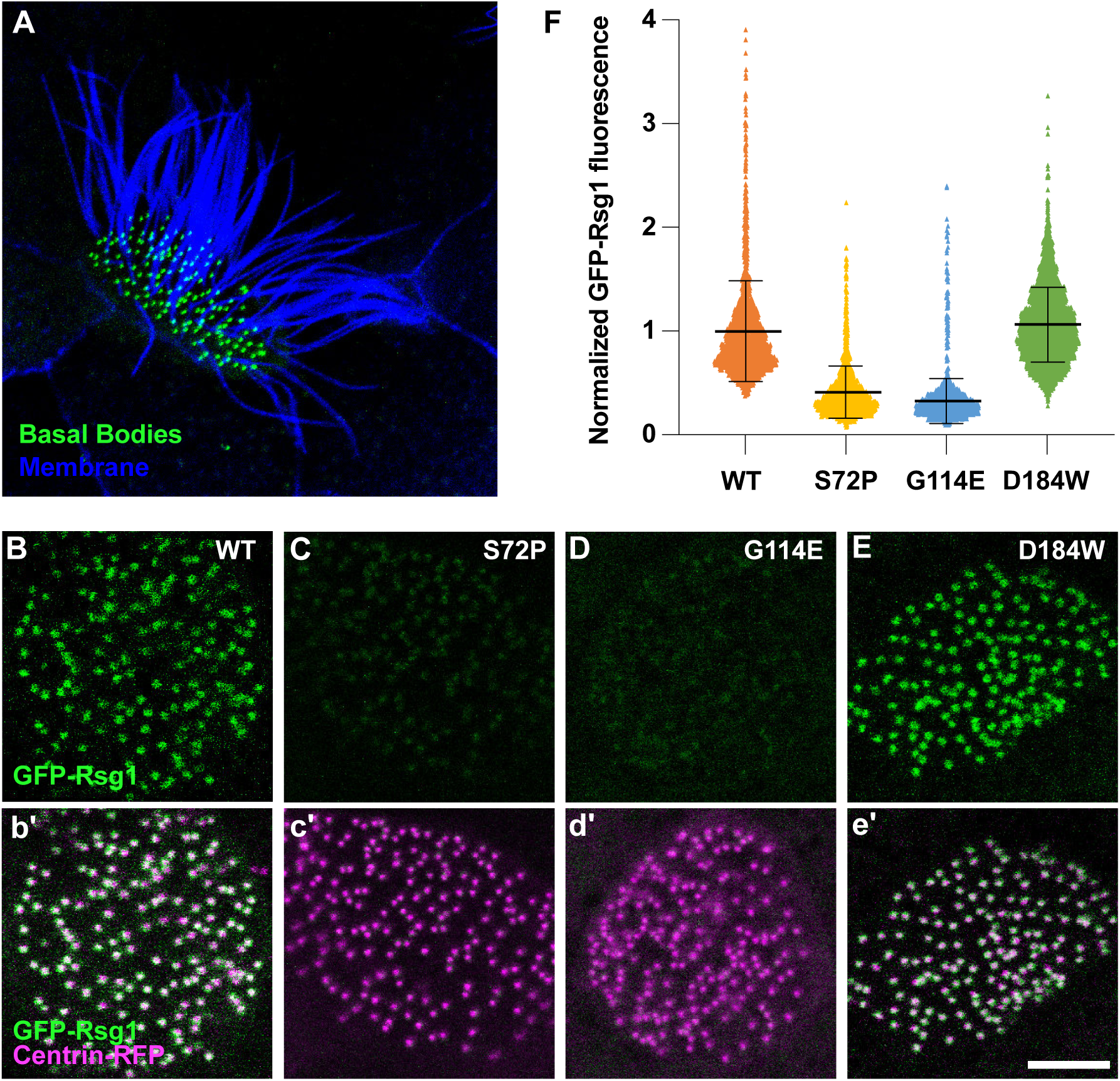
Ciliopathy-associated alleles of *RSG1* elicit distinct effects. (A) En face *in vivo* imaging of a *Xenopus* multiciliated cell with axonemes labelled by membrane-RFP (blue) and basal bodies labeled with centrin-BFP (green). (B-E) En face images of single MCCs showing Rsg1 basal body localization for control, and S72P (= human A76P), G114E (= human G118E), and D184W (= human R188W) variants. (b’-e’) merged channels showing Rsg1(green) with centrin (magenta), scale bar= 5um. (F) Graph showing mean ± standard deviation of normalized GFP-Rsg1 basal body fluorescence (see Methods). N> 25 cells in 5 embryos across 3 experiments for all conditions. N values and statistic can be found in Supp. Table 2.

Interestingly, the D184W (= R188W) variant displayed essentially normal recruitment to basal bodies (**Fig. 2E**, **F**) and also generated normal levels of protein in western blots (**Supp. Fig. 4**). These data suggest that G118E pathogenicity stems largely from a failure to be recruited to basal bodies, while the pathogenesis of R188W stems not from a change in its localization, but from a change in its function. We therefore sought to further test the impact of these ciliopathy alleles *in vivo*.

### Ciliopathy-associated alleles of *CPLANE2/RSG1* disrupt IFT-A2 recruitment to the base of cilia

CPLANE is implicated in the recruitment of IFT-A2 to basal bodies and ciliopathy-associated alleles of *JBTS17* and *INTU* disrupt this function ^11,12,22^. As a direct test of pathogenicity, we asked if ciliopathy alleles of *CPLANE2/RSG1* did likewise. First, we confirmed that Rsg1 knockdown (KD) disrupted BB recruitment of the IFT-A2 subunit Ift43 ^12^, and that this defect was significantly rescued by co-expression of wild-type Rsg1 (**Fig. 3A-C, G**). Consistent with the absence of a stable protein, co-expression of S72P did not significantly increase recruitment of Ift43-GFP over the knockdown (**Fig. 3D, G**). Likewise, and consistent with the failure of G114E to localize to basal bodies BB’s, expression of this allele also failed to significantly increase Ift43 levels above the knockdown (**Fig. 3E, G**).

**Fig 3.**
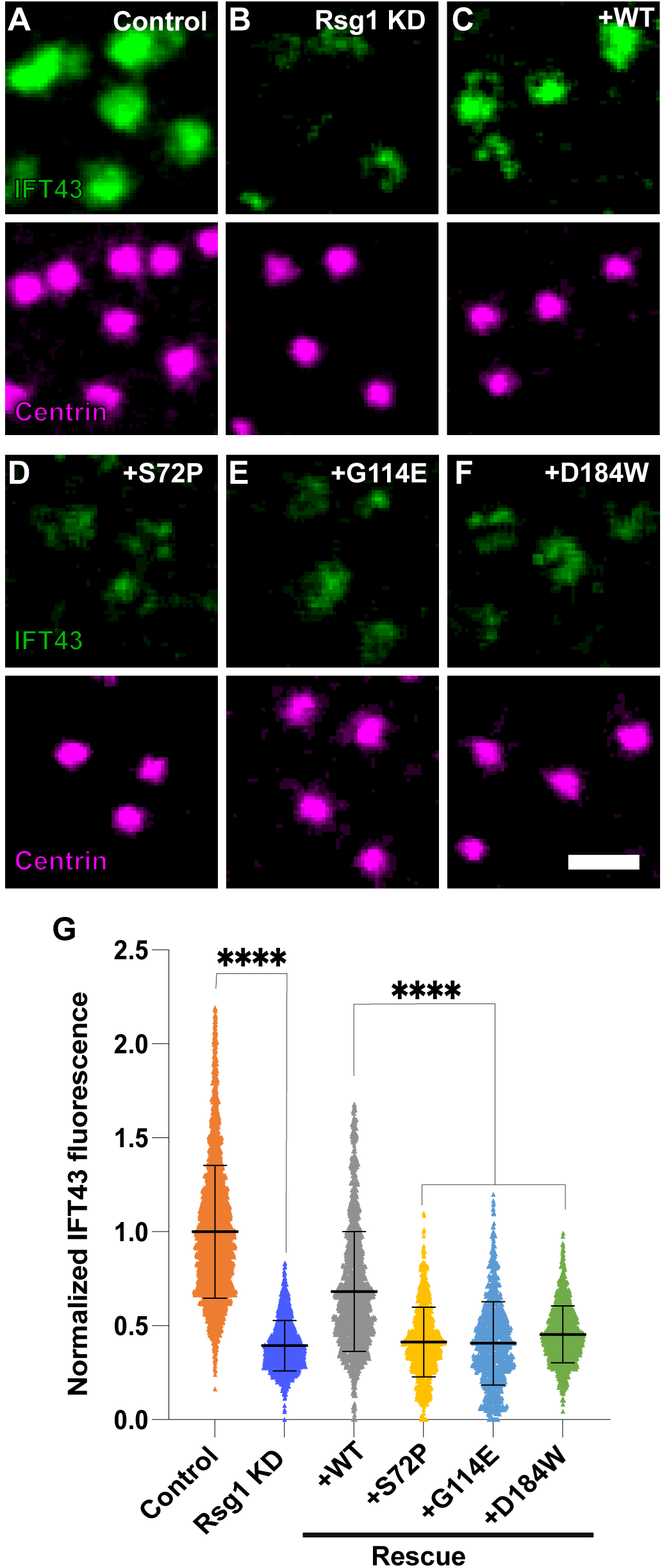
Ciliopathy-associated alleles of *RSG1* disrupt IFT-A2 recruitment to the base of cilia. (A-F) High magnification *en face* images showing IFT43 (green) localization at basal bodies labeled by centrin (magenta). (A-C) Control, Rsg1 KD and rescue of KD with WT Rsg1. (D-F) Failure of rescue by indicated variant alleles. Scale bar= 1μm. (G) Graph showing mean ± standard deviation of normalized IFT43-GFP basal body fluorescence. N> 21 cells in 6 embryos across 2 experiments for all conditions. N values and statistics can be found in Supp. Table 2.

By contrast, the normally localized D184W did elicit a modest but significant increase in Ift43 levels as compared to knockdown (**Fig. 3F, G**). These levels were also significantly increased compared to co-expression of either S72P or G114E, suggesting that D184W is by comparison a quantitatively milder variant. Just the same, Ift43 levels after D184W expression remained significantly below control levels and below the levels of WT Rsg1 rescue (**Fig. 3G**). Thus, we conclude that all three ciliopathy alleles of *RSG1* are pathogenic at least in part because of their failure to mediate IFT-A2 recruitment to basal bodies.

### Ciliopathy-associated alleles of *RSG1* and *JBTS17* disrupt basal body docking

Loss of CPLANE subunits causes basal body docking defects in *Xenopus* and mice ^10,12,21^, but whether disruption of basal body docking contributes to CPLANE-associated ciliopathy remains unknown. We therefore asked if ciliopathy variants of *CPLANE2* impact basal body docking in *Xenopus* MCCs (**Fig. 4A**). To this end, we first confirmed that Rsg1 KD disrupted BB docking ^12^ and that this defect could be significantly rescued by co-expression of wild-type Rsg1 (**Fig. 4B-D, H**). As expected, S72P failed to rescue BB docking; mean BB depth after expression of this allele was not significantly different from knockdown (**Fig. 4E, H**). Interestingly however, G114E did rescue docking with modest significance and notably, D184W elicited a highly significant rescue of BB depth compared to knockdown (**Fig. 4F-H, H**). However, these rescues fell well short of normal, with BB depths remaining significantly different from controls and from rescue with WT Rsg1.

**Fig 4.**
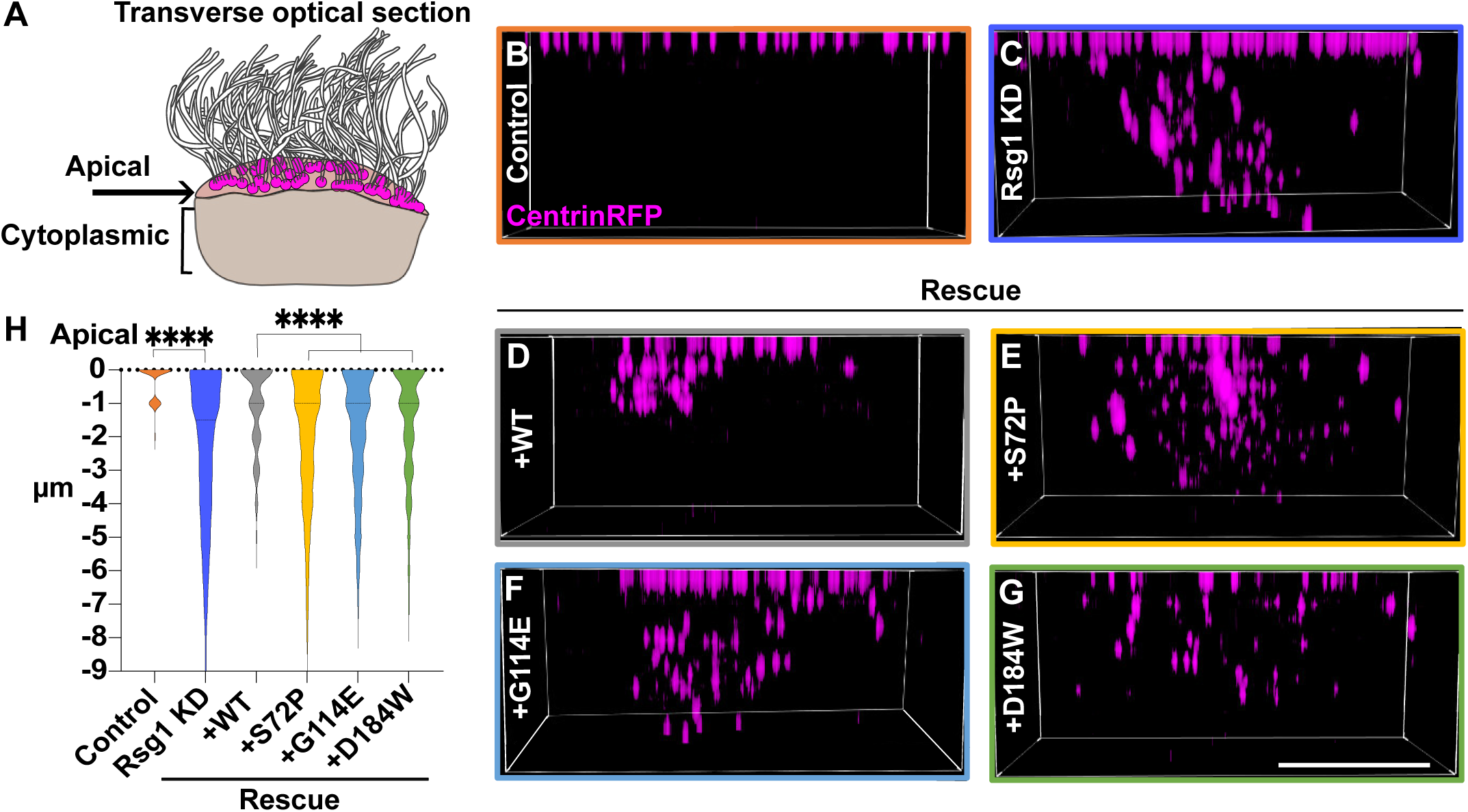
Ciliopathy-associated alleles of *RSG1* disrupt basal body docking. (A) Schematic representation of a MCC, depicting the apical and cytoplasmic regions shown in transverse optical section in this figure. (B-G) Transverse 3D projection of centrin (magenta), scale bar= 5um. (H) Graph shows the distribution of basal body depth below the apical surface in μm. N> 18 cells in 6 embryos across 2 experiments for all conditions. N values and statistics can be found in Supp. Table 2.

These results prompted us to ask if pathogenic variants in other CPLANE subunits may also act in part via BB docking. We found that knockdown of Jbts17 elicited severe basal body docking defects in *Xenopus* MCCs, and these could be effectively rescued by co-expression of wild-type Jbts17-GFP (**Supp. Fig. 5**). By contrast, expression of the ciliopathy-associated truncation Jbts17-Arg1569* ^36^ failed to rescue BB docking (**Supp. Fig. 5**). Thus, BB docking defects likely contribute to both RSG1-associated ciliopathy, and CPLANE-associated ciliopathy generally.

### GTP binding is required for Rsg1 association with CPLANE

Having found that pathogenic variants in *CPLANE2* cause human ciliopathy, we sought a deeper understanding of the encoded protein’s molecular functions. *RSG1* encodes a small GTPase. It binds GTP *in vitro* ^15^ and we previously showed that the canonical T→N mutation in the GTP-binding pocket disrupts basal body localization and imparts dominant-negative activity *in vivo* ^14^. But while the pathogenic G118E allele lies squarely within the GTP binding pocket (**Fig. 1D**), the molecular consequences of GTP binding by Rsg1 have never been explored.

We therefore used affinity purification and mass spectrometry (APMS) in ciliated mouse IMCD3 cells to compare the interactomes of wild-type Rsg1 and GDP-locked Rsg1^T69N^. Consistent with previous studies ^15,22^, APMS with wild-type Rsg1 strongly enriched all known CPLANE components, including not only Fuz, Intu, and Wdpcp, but also Jbts17 (**Table 1; Supp. Table 3**). Strikingly, interactions with all four CPLANE subunits were almost completely absent for Rsg1^T69N^ (**Table 1; Supp. Table 3**), suggesting the interactions are GTP-dependent. This result strongly suggests that GTP loading of Rsg1 is essential for its function and that other relevant interaction partners might also be identified by their specific failure to interact with Rsg1^T69N^. We therefore examined our interactomes for further insight into Rsg1 function.

**Table 1.** APMS comparison of WT and T69N Rsg1. Table shows spectral counts for selected proteins in APMS with WT Rsg1 versus the Rsg1(T69N) mutant. Interactors are grouped by function. Examples of non-specific interactors are provided at the bottom. Full data table can be found in Supp. Table 3.

|  |  | **Rsg1** | **Rsg1<sup>T65N</sup>** | **Log2FC** |
| --- | --- | --- | --- | --- |
| Cplane | **Fuz** | **1024** | **0** | **-11.19** |
|  | **Intu** | **430** | **0** | **-9.21** |
|  | **Wdpcp** | **1696** | **8** | **-8.51** |
|  | **Jbts17** | **898** | **0** | **-8.03** |
| Vesicle trafficking | **Hps1** | **208** | **0** | **-8.55** |
|  | **Rab3b** | **32** | **0** | **-7.08** |
|  | **Rab5c** | **22** | **0** | **-6.49** |
|  | **Rab23** | **10** | **0** | **-5.7** |
| Actin regulator | **Pf1** | **12** | **0** | **-6.31** |
|  | **Pdlim4** | **18** | **0** | **-5.86** |
|  | **Coro1c** | **20** | **0** | **-5.24** |
| Transition Zone | **Fam92a** | **38** | **0** | **-7.03** |
|  | **Dzip1** | **10** | **0** | **-5.98** |
|  | **Cby1** | **20** | **0** | **-4.44** |
| Non-specific | **Ruvbl1** | **128** | **100** | **-0.13** |
|  | **Eef1a1** | **448** | **472** | **0.3** |
|  | **Tubb6** | **1890** | **2756** | **0.77** |
|  | **Hsp90ab1** | **268** | **226** | **-0.02** |

### The GTP-dependent Rsg1 interactome links CPLANE to vesicle transport, actin assembly, and the ciliary transition zone

Because Intu and Fuz form a hexa-longin GEF ^13,15^, it is interesting that apart from other CPLANE subunits the most robustly enriched protein in our APMS was Hps1 (**Table 1; Supp. Table 3**), a subunit of the BLOC3 hexa-longin GEF complex ^37^. Also enriched was Rab5c (**Table 1; Supp. Table 3**), which is a known substrate of yet another hexa-longin GEF, Mon1-Ccz1-Bulli ^38^. These interactions warrant further investigation, as they raise the possibility of promiscuity, not just among effectors and substrates, but also among subunits of hexa-longin domain GEFs. In addition, several other Rabs were enriched specifically by wild-type Rsg1 but not Rsg1^T69N^, including Rab3B and the ciliogenic Rab29 ^39^ (**Table 1; Supp. Table 3**).

CPLANE subunits have also been implicated in actin assembly ^9,40,41^. It was interesting, then, that several actin-binding proteins were enriched specifically by wild-type Rsg1 but not Rsg1^T69N^. The most interesting were Pfn1 and Pdlim4, which are both dysregulated in a mouse model of retinal ciliopathy ^42^, and Coro1C, which is present in the ciliary membrane proteome ^43^ (**Table 1; Supp. Table 3**).

Finally, we found GTP-dependent interaction between Rsg1, and three proteins related to the ciliary transition zone, Fam92a/Cibar1, Cby1, and Dzip1 (**Table 1; Supp. Table 3**). These three proteins interact with one another, are essential for ciliogenesis, and are implicated in human ciliopathy ^44–49^. Because the CPLANE proteins have not been associated previously with the ciliary transition zone, we explored these interactions further.

### Modeling predicts Rsg1 interacts directly with the C-terminus of Fam92a

To understand the nature of the RSG1 interaction with the TZ, we used AlphaFold3 (AFold3) to explore its structural basis ^33^. Only FAM92A was predicted to interact directly with Rsg1 (**Fig. 5**), though the known interactions among FAM92A, CBY1 and DZIP1 ^45,47,50^ were each also predicted by Afold (not shown). Fam92a is a BAR domain protein that exists predominantly in a dimeric form ^51^ and Rsg1 forms a stable complex with Fuz ^15^, so we modeled a FAM92A homodimer with two RSG1/FUZ heterodimers (**Fig. 5B, C**). The FAM92A BAR domains formed a strongly predicted curved dimer with one RSG1/FUZ dimer predicted to bind at each end (**Fig. 5B, C**). This structure was consistently predicted across multiple seeds with Afold3 (not shown) and also predicted by AFold2 multimer ^52^ (**Supp Fig. 4).**

**Fig 5.**
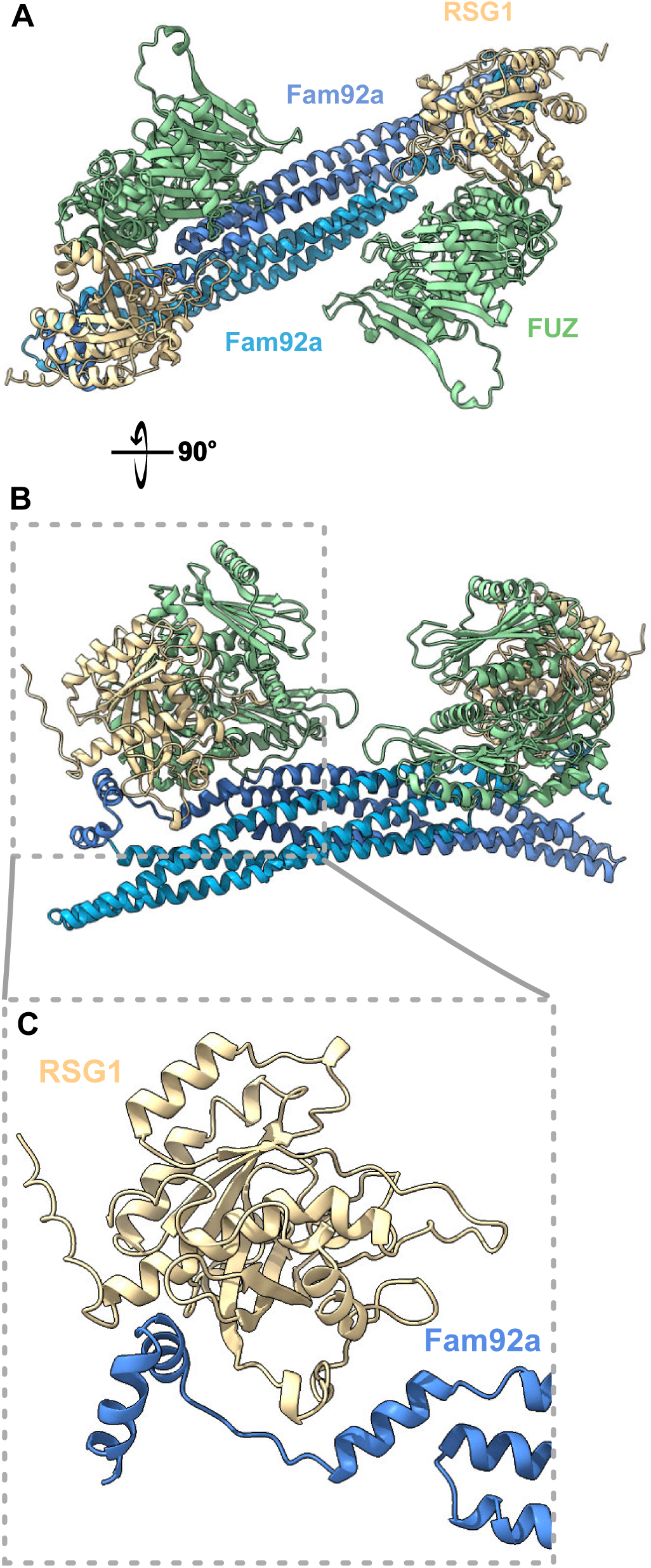
AlphaFold3 predicts direct interaction of RSG1 with FAM92A. (A) Alphafold3 prediction of the structure of two human FUZ-RSG1 heterodimers interacting with one Fam92a homodimer. Colors indicate monomers as indicated. (B) 90-degree rotation from B. (C) Increased magnification view of C showing the RGS1-FAM92A interaction (after removal of FUZ and one copy of FAM92A for clarity).

We performed several additional tests of the veracity of this prediction. First, we modeled interaction of monomers of RSG1 and FAM92A, and the Predicted Aligned Error (PAE) suggested that the structure of each monomer was confidently modeled, as was the interface of the key C-terminal helix of FAM92 with RSG1 (**Fig. 6A**, arrows). The interface predicted template modeling (ipTM) score for the overall model was 0.78 and the predicted local distance difference test (pLDDT) score for the C-terminal helix of FAM92A that contacts RSG1 was >70 (**Fig. 6B**). Even stronger scores were found when we modeled and RSG1 monomer with only the C-terminus of FAM92A (**Fig. 6C, D**). By contrast, no interaction was predicted when queried with RSG1 and FAM92A lacking the C-terminus (**Fig 6E, F**).

**Fig 6.**
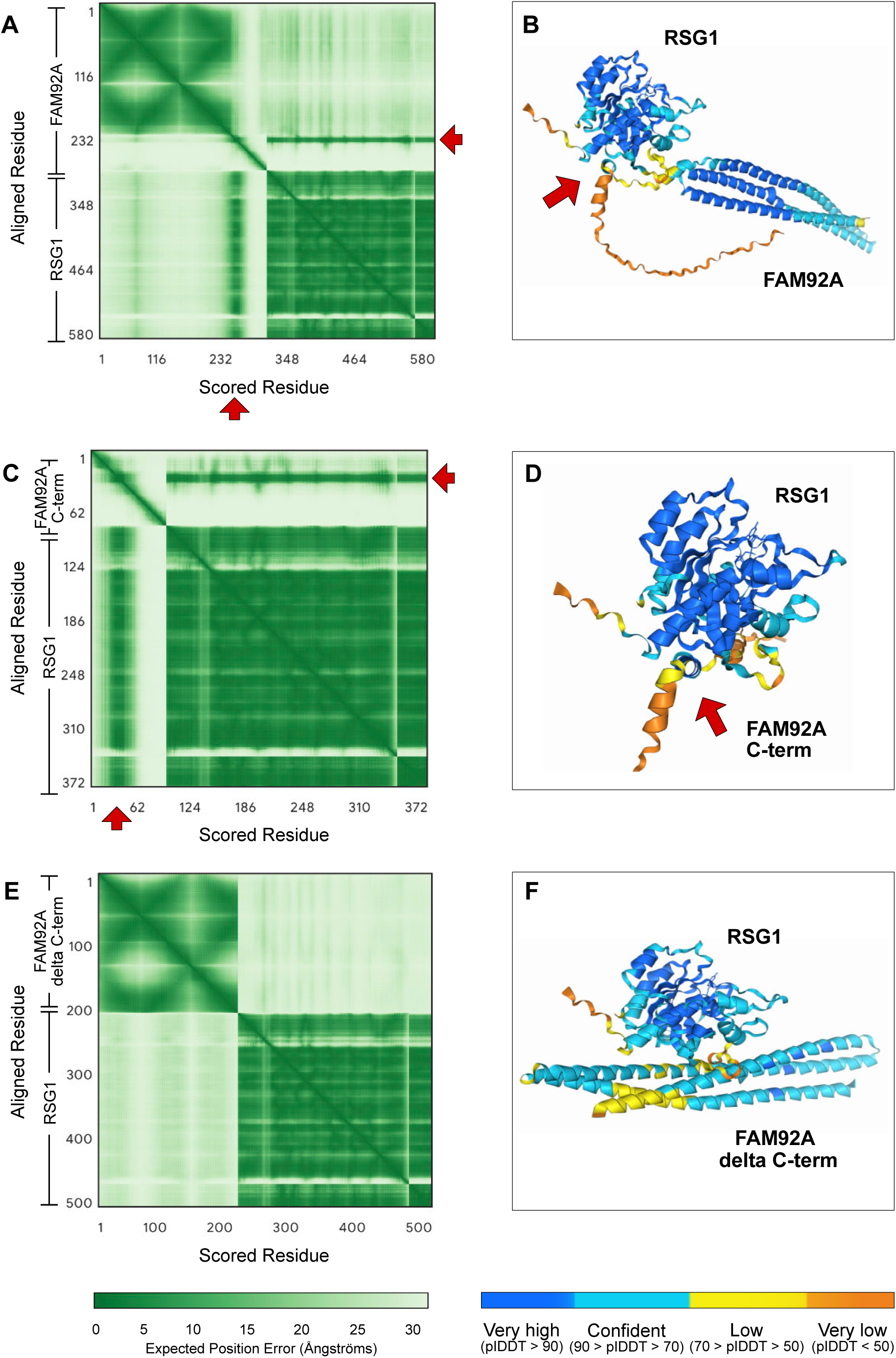
Quantification of AlphaFold3 Structure Predictions. (A) PAE Plot for the interaction of full-length human RSG1 and FAM92A monomers; arrow indicates region of Rsg1 interaction with the C-terminus of Fam92. (B) Structure of RSG1 monomer in complex with FAM92A monomer, with colors indicating PlDDT. (C) PAE Plot for the interaction of full-length RSG1 with the C-terminal 83 amino acids of FAM92A; arrow indicates region of Rsg1 interaction. (D) Structure corresponding to C, with colors indicating PlDDT. (E) PAE Plot indicates no interaction between full-length RSG1 with the C-terminal 83 amino acids of FAM92A; arrow indicates region of Rsg1 interaction. (F) Structure corresponding to E, note very poor PlDDT scores at the region of “interface.”

This overall structure of Fam92 interaction with Rsg1 calls to mind the paired dimer structure of Rab GTPases bound to their effectors, such as Rab10 and Mical1 or Rab11b and FIP2 ^53^. Moreover, the combined involvement of the N-terminus, the α3-β5 loop, and the C-terminal region of the α5 helix of Rsg1 to interact with the C-terminal helixes of FAM92A (**Fig. 6D; Supp. Fig. 2**) bears at least some similarity to the mechanism revealed by crystallography for interaction of Rab3a with the C-terminus of its effector Rabphilin3 ^54^.

Notably, a ciliopathy-associated C-terminal truncation of FAM92A fails to localize to basal bodies and instead accumulates in the ciliary tip ^44^. To link that finding to our work here, we made the corresponding truncation of *Xenopus* Fam92a, and it also displayed reduced basal body localization and ectopic accumulation in axonemes (not shown). This effect resembles that seen for another transition zone protein, Cep162, which also moves ectopically to ciliary tips when not tethered to basal bodies ^55^.

### RSG1 is required for ciliogenesis and recruitment of FAM92A to basal bodies in human cells

Rsg1 is essential for ciliogenesis in *Xenopus* and mice ^14,21^, but neither its role in human cells nor its relationship to Fam92 have yet been tested. We therefore used CRISPR to generate two distinct *CPLANE2/RSG1* knockout lines in human RPE1 cells (**Supp. Fig. 7A, B**). In both lines, Rsg1 loss resulted in roughly 75% reduction of ciliation as indicated by immunostaining for Arl13b (**Fig. 7A-C, G**). Moreover, FAM92A recruitment to basal bodies was severely, if not completely, reduced in each of the *RSG1* KO lines (**Fig. 7D-F, H**). This result was specific, as neither line displayed loss of the centrosomal protein Cep192 (**Fig. 7D-F**). These data suggest that the disease phenotypes we observed in patients with variants in *RSG1* (**Fig. 1**) indeed result from loss of cilia and that they are related at least in part to loss of FAM92a from basal bodies.

**Fig 7.**
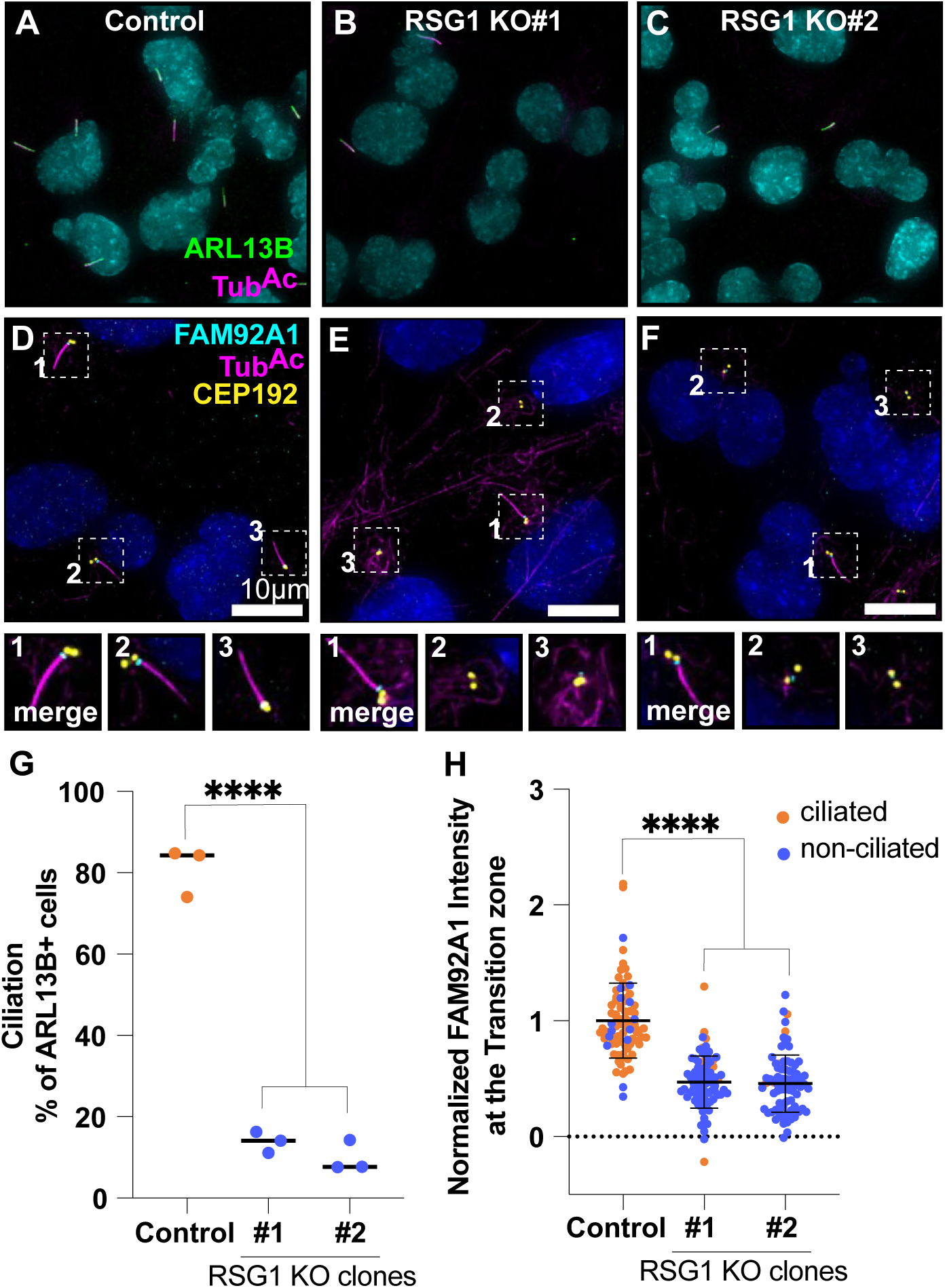
Rsg1 is required for ciliogenesis and recruitment of Fam92a to basal bodies in human cells. (A-C) Human RPE1 cells stained for ARL13B (green) and TubAc (magenta) as markers in control and Rsg1 crispant cells. scale bar=10µm (D-F) Imaging of Fam92a (cyan) fluorescence at the transition zone CEP192 (yellow) in RPE1 cells with mutated Rsg1. Insets show three indicated cilia from the panels above. (G) Percentage of ARL13B (green) positive cells in Rsg1 mutants. (H) Mean ± standard deviation of normalized Fam92a at the transition zone, ciliated cells marked in orange and non-ciliated in blue. N> 80 for all conditions. N values and statistics can be found in Supp. Table 2.

### CPLANE proteins are required for normal transition zone protein recruitment

Fam92a is present at mammalian ciliary transition zone (TZ), but its role at this location has been well-defined only in *Drosophila* ^44,45 48^. The loss of FAM92a from basal bodies after RSG1 loss therefore prompted us to ask if the CPLANE complex may be more generally involved in TZ architecture in mammalian cells. NPHP1 is a canonical TZ marker and a well-known ciliopathy protein ^56^, and we found that NPHP1 was severely reduced from basal bodies after RSG1 loss in human RPE1 cells (**Fig. 8A-C, G**), though like Fam92, it was not entirely eliminated.

**Fig 8.**
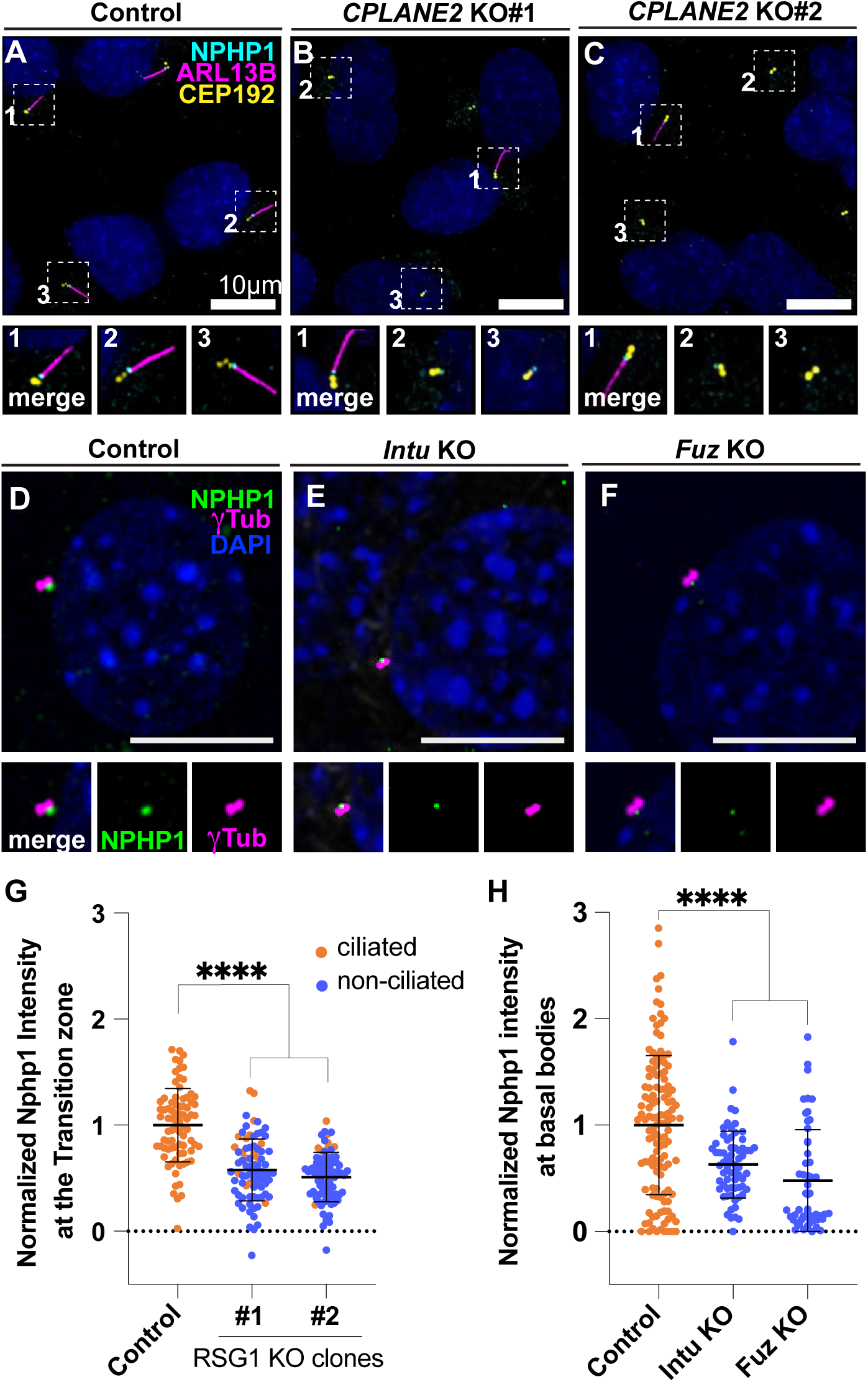
CPLANE is required for normal recruitment of the transition zone to the basal body. (A-C) Human RPE1 cells stained for the TZ protein Nphp1, the basal body protein Cep192, and the cilia marker Arl13b. Scale bar= 10μm. (G) Graph shows mean ± standard deviation of normalized Nphp1 fluorescence at the transition zone for indicated genotypes. (D-F) Nphp1(green) fluorescence at basal bodies labeled in tub(magenta) in *Intu or Fuz* mutant mouse embryo fibroblasts. Scale bar = 10μm. (H) Graph shows mean ± standard deviation of normalized Nphp1 fluorescence at the basal body in controls and *Intu* or Fuz mutants. N> 52 for all conditions. N values and statistics can be found in Supp. Table 2.

Finally, we were curious to know if this effect on the TZ was specific to loss of Rsg1, or if it may be a general feature of the larger CPLANE complex. We therefore examined mouse embryonic fibroblast cell lines that were lacking either Fuz or Intu, two other CPLANE subunits ^14,17^. Both lines displayed severe defects in ciliogenesis as expected (**Supp. Fig. 7C-G**), and both also displayed significant, but not total, loss of Nphp1 from basal bodies (**Fig. 8D-F,H**). Thus, disruption of normal TZ protein recruitment to basal bodies is a common feature of CPLANE loss and may be related to CPLANE-associated ciliopathy.

### Conclusion

Here, we have shown that variants in the *CPLANE2/RSG1* gene cause human ciliopathy, and these variants severely disrupt either the localization or the function of Rsg1. We further show that disruption of GTP binding inhibits Rsg1 interaction with other CPLANE components as well as with novel interactors. AlphaFold predicts that one of those, Fam92a, is a novel effector that is recruited to basal bodies by the Rsg1 GTPase. Finally, these results led us to discover that recruitment of transition zone proteins to basal bodies is a shared feature of multiple CPLANE subunits. These data are significant for several reasons.

First, these data shed light on the still enigmatic mechanism of Rsg1 function. Though similar to Rabs, Rsg1 displays many notable differences such as very low catalysis of GTP due to lack of the crucial glutamine in the G3 region ^15^. That said, our data strongly suggest that GTP loading is essential, both for CPLANE interaction and for binding to Fam92a (**Table 1**). Moreover, Rabs can simultaneously bind multiple effectors ^53^, and our data suggest the same is true for Rsg1. Indeed, our APMS and modeling suggest that Fam92a is an Rsg1 effector protein that can bind simultaneously with the known effector Fuz (**Fig. 5A-D**).

Second, CPLANE is known to affect both retrograde IFT and basal body docking in *Xenopus* ^10–12^, but only the former had previously been implicated in disease etiology ^22^. It is significant, that disease alleles of two CPLANE subunits fail to support basal body docking (**Fig. 4; Supp. Fig. 5**). Moreover, the interaction of Rsg1 with Fam92, Cby1 and Dzip1, is notable because like CPLANE proteins, these are also implicated in basal docking and have also been associated with defects in retrograde IFT ^47–49^. Further exploration of these proteins’ functional interaction will be of interest.

Third, the identification of Fam92 as a CPLANE interactor provides new insight into the complex interplay of proteins and membrane at the base of cilia ^57^. Interestingly, the CPLANE subunits Fuz, Intu, and Wdpcp, but not Rsg1 can directly interact with lipids, especially PI(3)P, PI(4)P, and PI(5)P ^15^. Rsg1 does not bind directly to lipids, so it is interesting that it does bind directly to Fam92, whose BAR domains bind and tubulate negatively charged lipids in vitro ^51^. Moreover, Fam92 interacts with lipids via a series of lysine and arginines on the concave face of the dimer ^51^, while Rsg1 and Fuz are positioned opposite, on the convex side (**Supp. Fig. 6C**). Thus, Rsg1 could interact with lipid-engaged Fam92, providing a possible route to recruitment of CPLANE to curved membrane surfaces. These findings complement previous work on BAR-domain proteins in ciliogenesis, such as MiniBAR and Pacsins ^58,59^.

Finally, these data provide insights into the molecular basis of the wide range of multi-organ congenital disorders termed ciliopathies ^60^, including OFD, a genetically heterogeneous ciliopathy characterized by anomalies of the oral cavity, face, and digits. Affected individuals may also present additional malformations including cerebral and/or cerebellar structural anomalies, cystic renal disease, and skeletal and ocular anomalies ^61^. Our work describes three individuals with OFD phenotype from three unrelated families in whom four distinct variants in CPLANE2 were identified. These OFD-affected individuals showed no other variants in genes previously associated with this condition after exome sequencing. Individuals one and two of our series presented with cardinal orofacial and digital OFD features in addition to variable expressivity of other clinical phenotypes. In contrast, individual 3 presented with a milder phenotype including OFD manifestations but not all of the cardinal ones. Importantly, our functional studies are consistent with this observation, as the milder phenotype observed in participant 3 correlates with less severe disruption in IFT recruitment and basal body docking in comparison with individuals one and two. Thus, our work suggests RSG1 is an OFD locus, and this locus should be investigated in additional cases of OFD syndrome to allow a better delineation of the phenotype, follow-up and the appropriate genetic counselling to the families.

## Acknowledgements

This work was supported by: The NICHD (R01HD085901) to E.M.M. and J.B.W; R01AR054396 and R01HD089918 to J.F.R.; R01GM121565, P01CA254867, R01DK127665 and R03TR004209 to P.K.J.; NIH R35GM122480 and the Welch Foundation (F-1515) to E.M.M.; Support from the Spanish Ministry of Science and Innovation (MICINN) through the program Juan de la Cierva to N.M.-G.; Swiss National Science Foundation (SNSF) grant number 32003B_219808 to I.F.; Support from Instituto de Salud Carlos III-FIS project PI22/00287, Xunta de Galicia (Centro de Investigación de Galicia CINBIO 2023-2027) Ref. ED431G-2023/06. To D.V.;An FPU predoctoral fellowship from the Spanish Ministry of Education, Culture and Sports (FPU 19/00175) to C. S.; and an ARCS Fellowship to C.D.

## Supplemental Materials and methods

### Patients

We detected three patients with a phenotype consistent with ciliopathy and with homozygous or compound heterozygous variants in *RSG1* from three di9erent centers. Clinical investigators were contacted through GeneMatcher, which enables new gene phenotype connections ^62^. All clinical and molecular data were collected including prenatal, morphologic, and neurodevelopmental characteristics as well as previous genetic studies. Informed consent approved by the local IRB was signed by their legal guardians.

### Patient Genetic analysis

Exome sequencing (ES) was performed for all patients according to the protocols and platforms of each center. Evaluation of ES variants excluded pathogenic changes in previously described OFD genes in any of the probands, which prompted us to search for a new gene responsible for this condition. In general, the identification of CPLANE2 variants in the sequencing data was done by filtering for: 1) variants with an ultra-rare allele frequency in the population (not present in gnomAD) and/or 2) variants located in exonic regions or splice sites. Interpretation of single nucleotide variants was done according to ACMG guidelines 4 ^63^.

### *Xenopus* embryo manipulations

Follow-up experiments on select candidates were performed in *Xenopus laevis*. All *Xenopus* experiments were conducted in accordance with the animal protocol AUP-2024-00130 and the animal ethics guidelines of the University of Texas at Austin.

Female adult *Xenopus laevis* were injected with human chorionic gonadotropin the night preceding experiments to induce ovulation. Females were squeezed to lay eggs, and eggs were fertilized in vitro with homogenized testis in 1/3X Marc’s modified Ringer’s (MMR). Two-cell stage embryos were de-jellied in 1/3X MMR with 2.5% (wt/vol) cysteine (pH 7.9), then washed and maintained in 1/3X MMR solution. For plasmid, mRNA microinjections, embryos were placed in 2% Ficoll in 1/3X MMR and injected using a glass needle, forceps, and an Oxford universal micromanipulator. After injection, embryos were kept in Ficoll for at least 30mins and removed from Ficoll solution to develop in 1/3 MMR solution.

### Plasmid, mRNA, and MO microinjections

Plasmids containing GFP-Ift43, GFP-Rsg1, or Centrin-RFP were used for mRNA synthesis^12^. To generate a *Xenopus* allele corresponding to the human patient allele, mutagenesis was performed on the GFP-tagged Xenopus Rsg1, using Q5 Site-Directed Mutagenesis Kit (NEB, Cat # E0554S). Capped mRNAs were synthesized using the mMESSAGE mMACHINE SP6 transcription kit (Thermo Fisher Scientific, Cat # AM1340). Translation-blocking Rsg1 morpholino (5′-GGCCCGTATCTCTGTAGTGCAGCAA–3′) (Gene Tools) has been previously described ^12 14^. mRNA and/or morpholino were injected into two ventral blastomeres at the four-cell stage to target the epidermis. mRNAs were injected at 40∼100 pg per blastomere, and morpholino was injected at 30 ng per blastomere.

### Immunoblotting

Embryos were lysed in Lysis buffer (ThermoFisher Scientific, Cat# 78501) containing protease inhibitors. The lysates were centrifuged to remove cell debris, and the supernatants were subjected to SDS-PAGE followed by immunoblotting using standard protocols. The antibodies used were as follows: anti-GFP antibody (Santa Cruz, Cat# sc-9996), HRP-conjugated goat anti mouse IgG (H+L) secondary antibody (ThermoFiser Scientific, Cat# 31430), beta actin monoclonal antibody (Proteintech, Cat # 6009-1).

### Live imaging and image analysis for *Xenopus*

For live imaging, *Xenopus* embryos (stage 25) were mounted between two coverslips and submerged in 0.01% benzocaine in 1/3X MMR and imaged on a Zeiss LSM700 confocal microscope using a Plan-Apochromat 63 × 1.4 NA oil DIC M27 immersion lens. Each experiment includes multiple biological replicates, conducted on different days. Data for Rsg1 allele localization experiments all include three replicates, Rsg1 rescue of IFT43 and docking phenotype data represent two replicates.

Images were processed using the Fiji distribution of ImageJ (Schindelin *et al.*, 2012) and figures were assembled in Affinity Designer.

Quantification of fluorescence intensity of basal body localizing proteins was performed on micrographs taken at the single z depth which best captured the most basal bodies as labeled by centrin-RFP. Quantification performed on single z micrographs for both basal body localization of both Rsg1 alleles and IFT43 rescue data.

Fluorescence intensity was measured by first duplicating the centrin-RFP channel and using the duplicate to reduce background signal as well as smoothing the image to then analyze particles. Measurements were then taken by using the particle analysis ROIs on the GFP-tagged candidate protein and centrin-RFP raw channels independently using the “Measure” function. Normalization was performed by calculating the ratio of candidate GFP intensity to the centrin-RFP intensity and normalizing all data sets to the control average.

### Tandem ABinity Purification

5 mL packed cell volume of hTERT RPE1 cells expressing LAPN-tagged proteins were re-suspended with 20 mL of LAP-resuspension bu9er (300 mM KCl, 50 mM HEPES-KOH [pH 7.4], 1 mM EGTA, 1 mM MgCl2, 10% glycerol, 0.5 mM DTT, protease inhibitor [A32965, Thermo Fisher Scientific]), lysed by gradually adding 600 μL 10% NP-40 to a final concentration of 0.3%, then incubated on ice for 10 min. The lysate was first centrifuged at 14,000 rpm (27,000 g) at 4°C for 10 min, and the resulting supernatant was centrifuged at 43,000 rpm (100,000 g) for 1 hr at 4°C to further clarify the lysate. High speed supernatant was mixed with 500 μl of GFP-coupled beads ^64^ and rotated for 1 hr at 4°C to capture GFP-tagged proteins, then washed five times with 1 mL LAP200N bu9er (200 mM KCl, 50 mM HEPES-KOH [pH 7.4], 1 mM EGTA, 1 mM MgCl2, 10% glycerol, 0.5 mM DTT, protease inhibitors, and 0.05% NP40). After re-suspending the beads with 1 mL LAP200N bu9er lacking DTT, and protease inhibitors, the GFP tag was cleaved by adding 40ug TEV protease and rotating tubes at 4°C for 16 hours. TEV-eluted supernatant was added to 100 uL of S-protein agarose (69704-3, EMD Millipore) to capture S-tagged protein. After washing three times with LAP200N bu9er lacking DTT and twice with LAP0 bu9er (50 mM HEPES-KOH [pH 7.4], 1 mM EGTA, 1 mM MgCl2, and 10% glycerol), purified protein complexes were eluted with 50 uL of 2X LDS bu9er and boiled at 95°C for 3 min. Samples were then run on Bolt Bis-Tris Plus Gels (NW04120BOX, Thermo Fisher Scientific) in Bolt MES SDS Running Bu9er (B0002, Thermo Fisher Scientific). Gels were fixed in 100 mL of fixing solution (50% methanol, 10% acetic acid in Optima LC/MS grade water [W6-4, Fisher Scientific]) at room temperature, and stained with Colloidal Blue Staining Kit (LC6025, Thermo Fisher Scientific). After the bu9er was replaced with Optima water, the bands were cut into eight pieces. The gel slices were then destained, reduced, and alkylated followed by in-gel digestion using (125 ng) Trypsin/LysC (V5073, Promega) as previously described ^65^. Tryptic peptides were extracted from the gel bands and dried in a speed vac. Prior to LC-MS, each sample was reconstituted in 0.1% formic acid, 2% acetonitrile, and water.

## Mass Spectrometry

Samples were run on Thermo Orbitrap Fusion at Stanford University Mass Spectrometry (SUMS).

### Data Analysis

Raw mass spec result files were processed using Bionic (Protein Metrics, Inc.) software (version: v2.16.11) with the following parameters: precursor mass tolerance: 20 ppm, fragment mass tolerance: 40 ppm; fragmentation type: QTOF/HCD; propionamide as Cys fixed modification, variable modifications of Met, His and Trp oxidation, Asn and Gln deamidation, and Ser, Thr and Tyr phosphorylation, with maximum 2 variable modification; trypsin with maximum 2 missed cleavages and fully specific mode; identifications were filtered at 0.01 FDR at the protein level; a FASTA library of all human refseq proteins (curated and predicted) was used that was downloaded on 2018/06/18. Spectral counts were converted to normalized spectral abundance factor (NSAF) values ^66^ and significance of enrichment of bait-association were calculated and protein interaction networks were constructed in Cytoscape, as previously described ^67^.

All proteomic data are available in the PRIDE database (Project accession: PXD055830; Project DOI: 10.6019/PXD055830).

### Protein structure modeling

Modeling was performed using AlphaFold3 ^33^.and AlphaFold2 ^52^ and computational alanine scanning was performed using BAlas ^68^.

### Human and mouse cell culture

RPE1 *RSG1* knockout cell lines were generated using two single guide RNAs synthesized by Synthego (sgRNA cut site 1: UUGCCCUAGGGCUGCUUGAG, sgRNA cut site 2: CCCACUCACCGGUGGUCUCG). Thermo Fisher Scientific Neon Transfection System was used for electroporation of sgRNAs with Truecut Cas9 v2. Monoclonal cell line DNA was isolated with QuickExtract DNA Extraction Solution, the cut site was amplified with PCR primers around the deletion site (Forward primer TTGTGTTAGGCCCCACTTCC, reverse primer GACCAAGGAGCCCATGTAGG) and Sanger sequenced to identify insertions or deletions at the cut sites. Monoclonal *RSG1* knockout cell lines were further validated with RT-qPCR to quantify fold change of *RSG1* expression (Forward primer ATCATATGCTGCTGGCTTGC, reverse primer TGGTCAAATTTGGAGCCGATG; forward primer spans exon-exon junction). RNA was isolated with Qiagen RNeasy Mini Kit, and cDNA was synthesized with BioRad iScript cDNA Synthesis Kit. PowerUp SYBR Green Master Mix was used to amplify the target and control cDNA sequences.

MEFs were grown in Advanced DMEM supplemented with 10% FBS, 1% GlutaMax, and 1% antibiotic-anti-mycotic.

### Immunostaining

RPE1 cells were grown on glass coverslips (Azer Scientific Inc, ES0117520) and fixed with either 4% PFA (Thomas Scientific, C993M24 or a combination of 1.5% PFA (diluted in 1x PBS) at RT for 4min, followed by ice-cold MeOH at −20°C for 4min. Cells were blocked in blocking solution (2.5% FBS, 200 mM glycine, 0.1% Triton X-100 in PBS) for 1 h at RT. Cells were then incubated in primary antibody diluted in the blocking solution for 1 h and rinsed with PBST (0.1% Triton X-100 in PBS), followed by secondary antibody staining prepared in blocking solution for 1 h and rinsed with PBST. DNA was stained with Hoechst 33342 (Thermo Fisher, H21492), and cells were mounted in ProLong Diamond Antifade Mountant (Fisher Scientific, P36970). Immunofluorescence images were acquired at room temperature (25°C) using a DeltaVision Elite (GE Healthcare) controlling a pco.edge sCMOS camera with near-perfect linearity across its 15-bit dynamic range. Images were acquired with a Plan Apochromat 60× 1.40 NA oil objective lens with 0.2-µm z-sections. All image acquisition was done in SoftWoRx (6.0; GE Healthcare).

MEFs were plated on glass coverslips, grown to confluence, and starved for 24h in OptiMEM. Cells were fixed for 10 min at RT with 4% PFA in PBS, followed by methanol for 3 min at −20°C. After fixation, cells were blocked for 1 h at RT or overnight at 4°C in dPBS + 0.1% Triton X-100 + 2.5% BSA (IF block). Cells were incubated with primary antibodies diluted in IF block overnight at 4°C. Coverslips were washed with dPBS, and then incubated with Alexa-conjugated secondary antibodies raised in goat or donkey (Invitrogen) at RT. After three dPBS washes, coverslips were mounted using ProLong Gold antifade (Invitrogen) and sealed with nail polish. Cells were imaged with a TCS SP5 microscope (Leica) or Zeiss LSM700 confocal.

### Antibodies

The following antibodies were used for RPE1 cells: rabbit-NPHP1 (1:200, Sigma-Aldrich, SAB1401267), rabbit-FAM92A1 (1:200, Thermo Fisher Scientific, 24803-1-AP), mouse-acetylated Tubulin (1:1000, Santa Cruz Biotechnology, sc-23950), goat-CEP192 (1:2000; gift from Andrew Holland’s lab, raised against CEP192 [amino acids 1–211]). Secondary antibodies conjugated to Alexa Fluor 488, 568 or 647 were used (1:1000, Life Technologies Corporation).

The following antibodies were used for MEFs: rabbit α-Nphp1 (gift of G. Pazour, University of Massachusetts Medical School, Worcester, MA; amino acids 1–209 [Benzing et al., 2001; Fliegauf et al., 2006]), rabbit α-Arl13b (Proteintech, cat# 17711-1-AP), goat α–γ-tubulin (Santa Cruz Biotechnology, Inc., cat# sc-7396), mouse α-detyrosinated tubulin (Millipore, cat# ab254154).

### Image analysis of NPHP1 & ciliogenesis

For quantitation of signal intensity at the centrosome in RPE1 cells, deconvolved 2D maximum intensity projections were saved as 16-bit TIFF images. Signal intensity was determined by drawing a circular region of interest (ROI) around the centriole (small ROI [ROI_S_]). A larger concentric circle (large ROI [ROIL]) was drawn around ROI_S_. ROI_S_ and ROI_L_ were transferred to the channel of interest, and the signal in ROI_S_ was calculated using the formula I_S_ − [(I_L_ − I_S_)/(A_L_ − A_S_) × A_S_], where A is area and I is integrated intensity.

FIJI (FIJI is Just ImageJ) was used for image analysis of MEFs. To quantify ciliogenesis, the total number of cilia and nuclei were manually counted, and % ciliation was calculated. For transition zone localization of NPHP1, a circle was used to measure the integrated density of NPHP1 and gTub for every basal body in the image. The background intensity was also calculated for each channel. For both NPHP1 and gTub, the background intensity was subtracted from the experimental intensity. Then, the corrected NPHP1 intensity was normalized to the corrected gTub intensity.

## Supplemental Figure Legends

**Supp. Fig 1.**
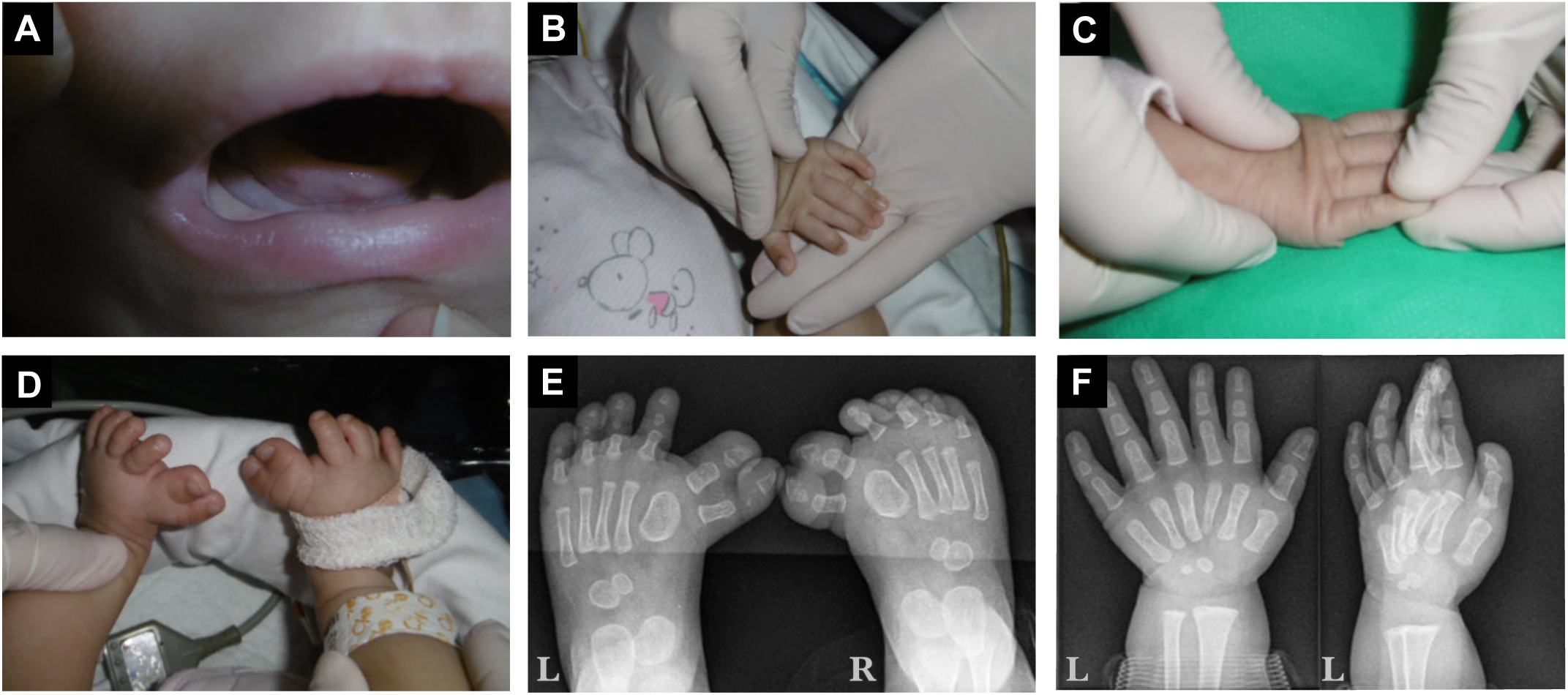
Photographs and X-rays of Patient 2. A. The image shows a lobulated tongue. B and C. Show polydactyly in both hands: insertional in the left hand and postaxial in the right hand. D. Preaxial polydactyly is observed in both feet. E and F. Plain X-rays of the same patient where polydactyly is evident in both hands and feet.

**Supp. Fig 2.**
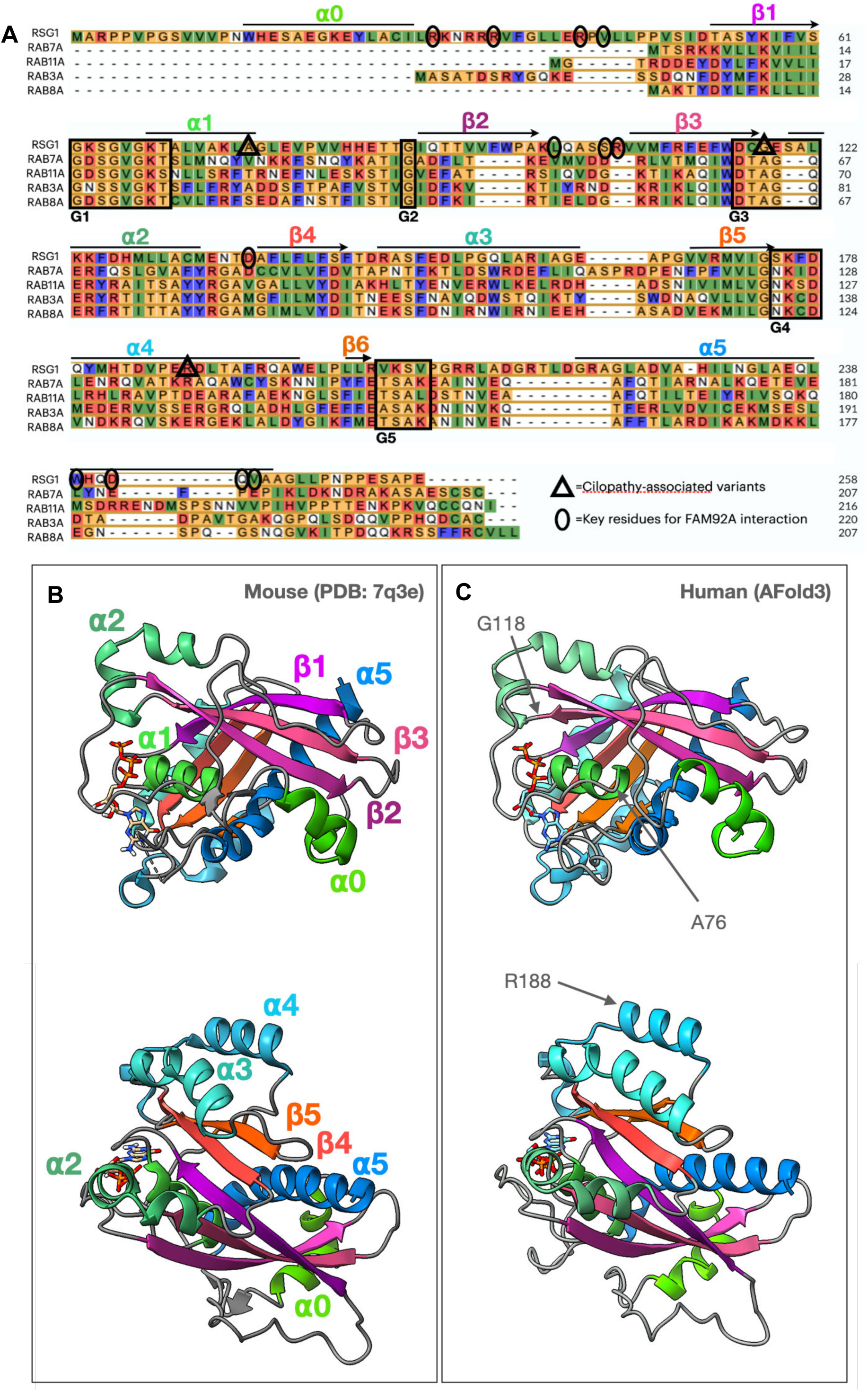
RSG1 sequence and protein structure. (A) Alignment of human RSG1 with related human RAB proteins. Triangles indicate ciliopathy loci identified here. Residues indicated by ovals are the top-scoring residues predicted to mediate interaction with RSG1 by computational alanine scanning using BAlas (see Methods). G domains are indicated by black boxes and structural elements related to Panel B are indicated above each sequenece. (B) Structure of Mouse Rsg1 (PDB: 7q3e) colored to indicate the conserved helicases and sheets characteristic of small GPTases. (C) Afold3-predicted structure of Human RSG1 colored to indicate the conserved helicases and sheets characteristic of small GPTases and the positions of ciliopathy loci identified here indicated in grey.

**Supp. Fig 3.**
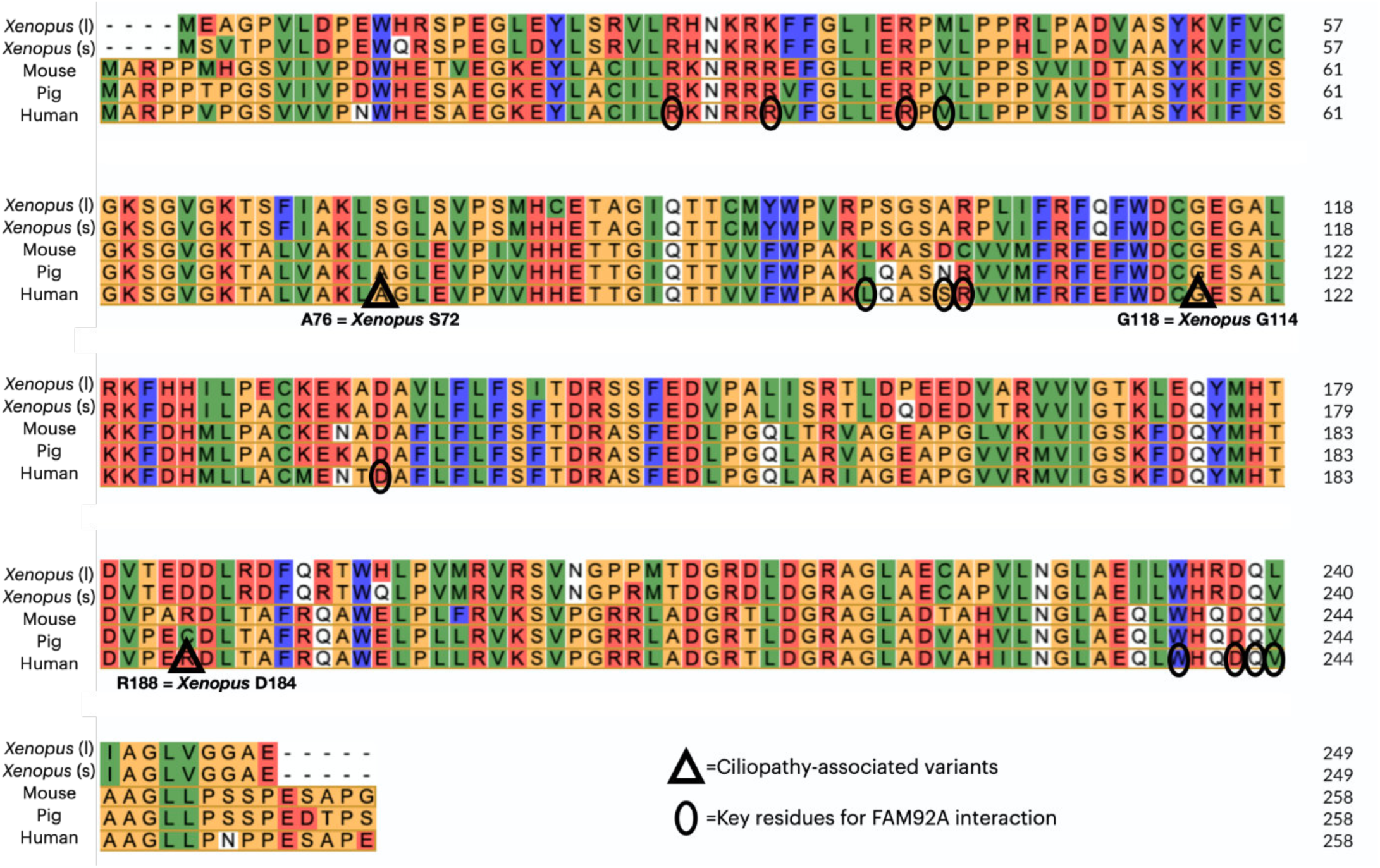
Alignment of vertebrate Rsg1 proteins. Triangles indicate ciliopathy loci identified here. Note high conservation of residues indicated by ovals, which are the top-scoring residues predicted to mediate interaction with RSG1 determined by computational alanine scanning using BALas (see Methods).

**Supp. Fig 4.**
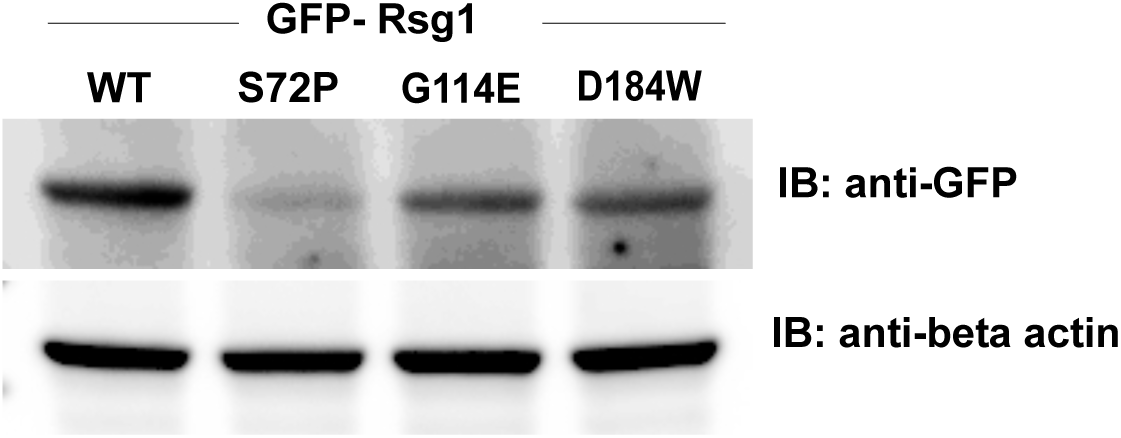
Protein stability of RSG1 disease alleles. Western blot showing protein levels for indicated disease alleles when expressed in *Xenopus* embryos.

**Supp. Fig 5.**
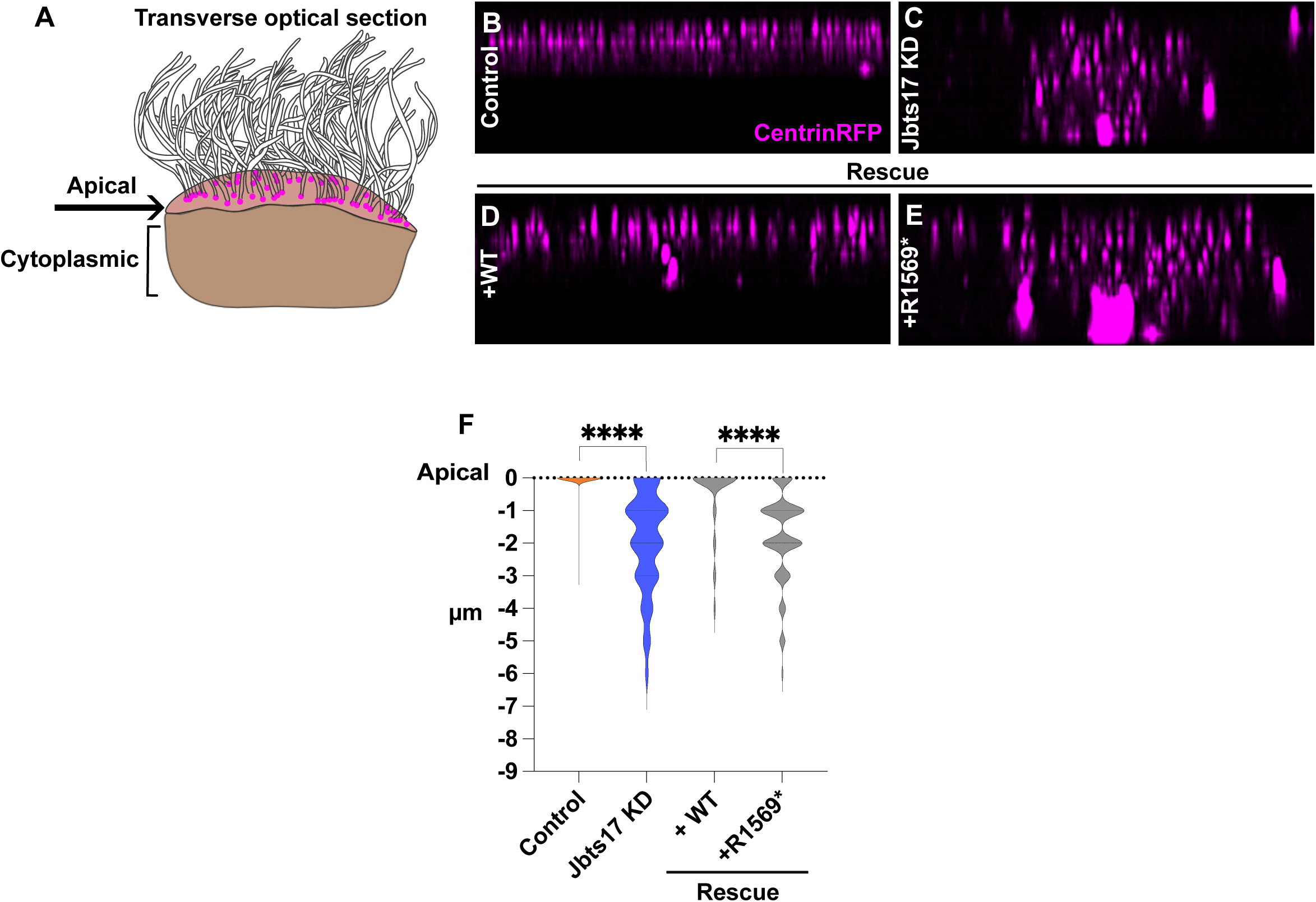
CPLANE Jbts17 disrupts basal body docking. (A) Schematic representation of a MCC, depicting the apical and cytoplasmic regions shown in the optical view. (B-E) Transverse 3D projection of centrin (magenta). (F) Graph shows the distribution of centrin signal through the cell at each μm depth starting from apical slice at 0. N> 12 cells for all conditions.

**Supp. Fig. 6.**
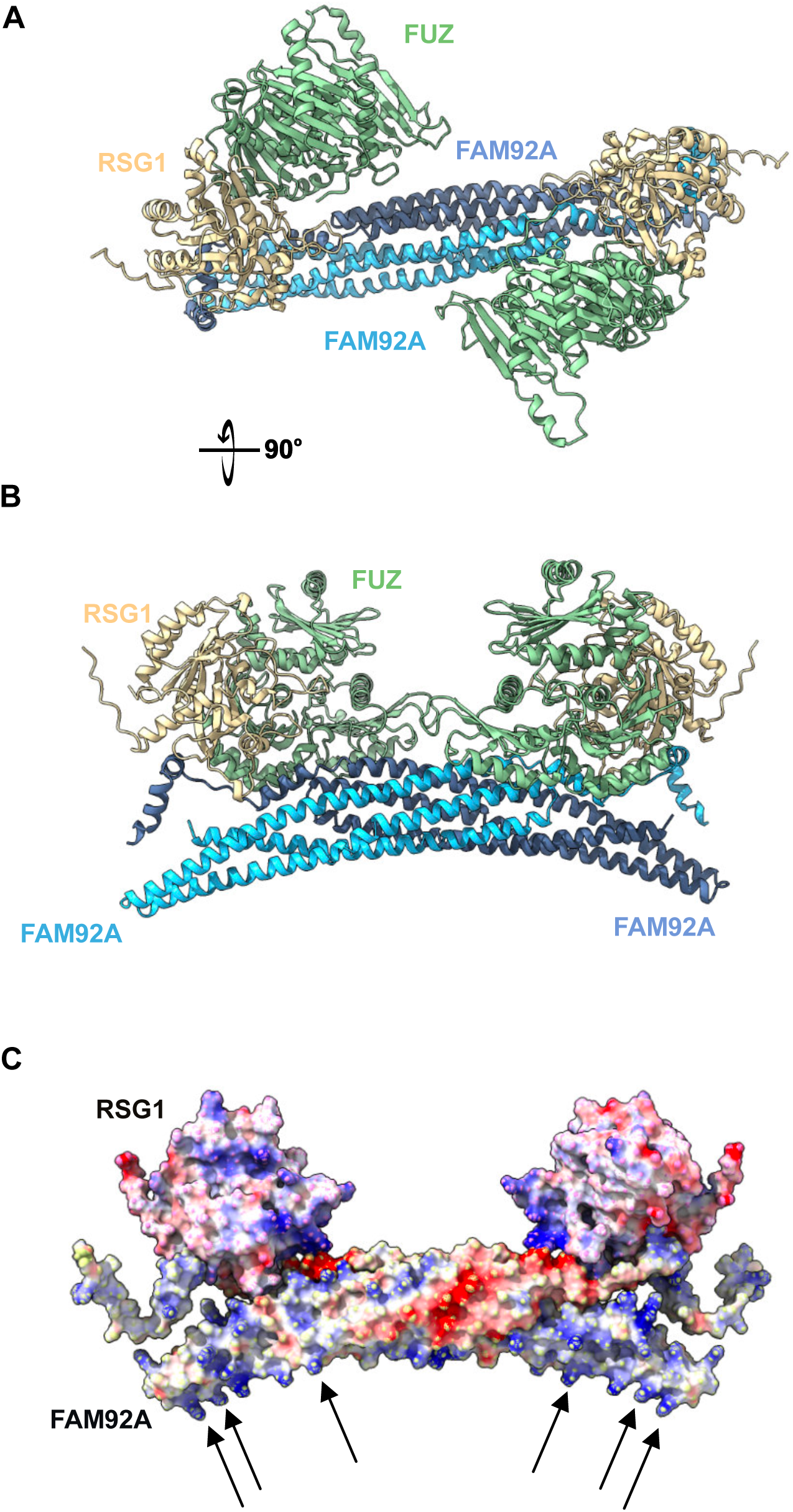
AlphaFold2 predicts direct binding between Rsg1 and the c-terminus of Fam92a. (A, B) AlphaFold2 model of FUZ/RSG1/FAM92A with a configuration similar to that predicted by Afold3 (compare with Fig. 6). (C) Figure shown in B with charges of residues indicated, arrows indicate lysines and arginines required for FAM92 binding to negatively charged lipids.

**Supp. Fig 7.**
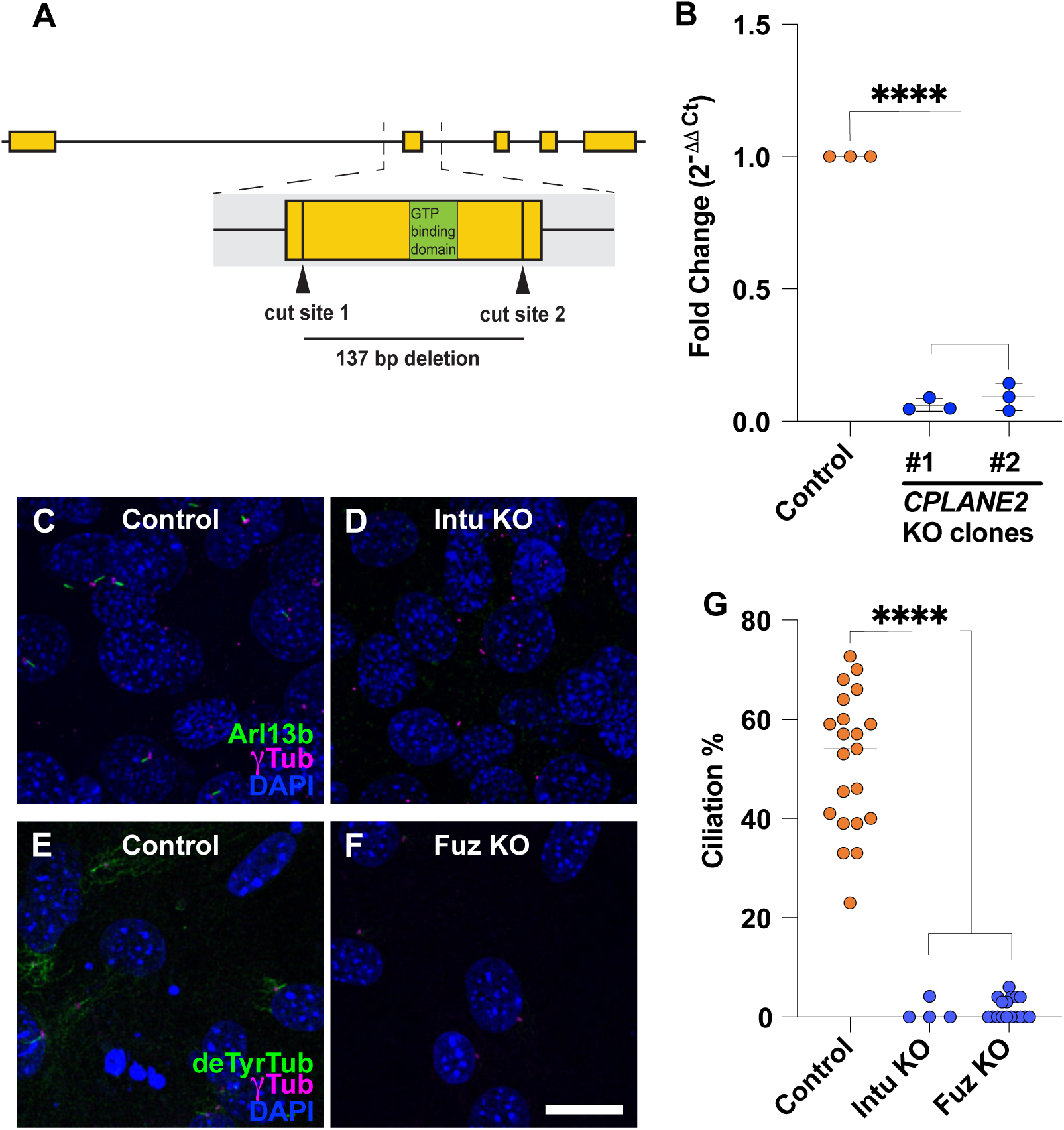
Generation and validation of RPE1 *RSG1* knockout cell lines and validation of Fuz and Intu mouse embryo fibroblasts. (A) CRISPR-Cas9 mediated *RSG1* knockouts were generated in RPE1 cells using two synthesized sgRNAs designed to cut on either side of the GTP binding domain within exon 2 of *RSG1*. Two monoclonal cell lines were isolated containing the same 137 base pair deletion. (B) RT-qPCR delta delta Ct analysis indicates a significant reduction in *RSG1* gene expression in both isolated knockout clones compared to a non-targeting control line. N=3 biological replicates, error bars are represented as mean with SD; **two-tailed P value ≤ 0.01 and ***two-tailed P value ≤ 0.001 (one-sample t test, theoretical mean = 1). (C-F) Immunofluorescence of mouse embryonic fibroblasts of indicated genotype with DAPI (blue) and gamma tubulin (magenta). Ciliation % was quantified using detyrosinated Tubulin(green) in Fuz and Arl13B (green) in Intu. Scale bar = 20μm. (G) Percentage of deTyrTub (green) and ARL13B (green) positive cells in *Fuz* and *Intu* mutants. N> 80. N values and statistics can be found in Supp. Table 2.

## Supplemental Table legends

**Supp. Table 1:** Spreadsheet of clinical data for patients.

**Supp. Table 2:** Statistics and N values for all experiments.

**Supp. Table 3:** Spreadsheet of proteomic data for APMS of WT Rsg1 and Rsg1(T69N). Full are available in the PRIDE database (Project accession: PXD055830; Project DOI: 10.6019/PXD055830).

## References

1 Hildebrandt, F., Benzing, T. & Katsanis, N. Ciliopathies. N Engl J Med 364, 1533–1543 (2011). 10.1056/NEJMra1010172

2 Reiter, J. F. & Leroux, M. R. Genes and molecular pathways underpinning ciliopathies. Nat Rev Mol Cell Biol 18, 533–547 (2017). 10.1038/nrm.2017.60

3 Mitchison, H. M. & Valente, E. M. Motile and non-motile cilia in human pathology: from function to phenotypes. J Pathol 241, 294–309 (2017). 10.1002/path.4843

4 Ishikawa, H. & Marshall, W. F. Intraflagellar Transport and Ciliary Dynamics. Cold Spring Harb Perspect Biol 9 (2017). 10.1101/cshperspect.a021998

5 Klena, N. & Pigino, G. Structural Biology of Cilia and Intraflagellar Transport. Annu Rev Cell Dev Biol 38, 103–123 (2022). 10.1146/annurev-cellbio-120219-034238

6 Tian, X., Zhao, H. & Zhou, J. Organization, functions, and mechanisms of the BBSome in development, ciliopathies, and beyond. Elife 12 (2023). 10.7554/eLife.87623

7 Reiter, J. F., Blacque, O. E. & Leroux, M. R. The base of the cilium: roles for transition fibres and the transition zone in ciliary formation, maintenance and compartmentalization. EMBO Rep 13, 608–618 (2012). embor201273 [pii] 10.1038/embor.2012.73

8 Adler, P. N. & Wallingford, J. B. From Planar Cell Polarity to Ciliogenesis and Back: The Curious Tale of the PPE and CPLANE proteins. Trends Cell Biol 27, 379–390 (2017). 10.1016/j.tcb.2016.12.001

9 Park, T. J., Haigo, S. L. & Wallingford, J. B. Ciliogenesis defects in embryos lacking inturned or fuzzy function are associated with failure of planar cell polarity and Hedgehog signaling. Nat Genet 38, 303–311 (2006).

10 Park, T. J., Mitchell, B. J., Abitua, P. B., Kintner, C. & Wallingford, J. B. Dishevelled controls apical docking and planar polarization of basal bodies in ciliated epithelial cells. Nat Genet 40, 871–879 (2008).

11 Brooks, E. R. & Wallingford, J. B. Control of vertebrate intraflagellar transport by the planar cell polarity eyector Fuz. J Cell Biol 198, 37–45 (2012). 10.1083/jcb.201204072

12 Brooks, E. R. & Wallingford, J. B. The Small GTPase Rsg1 is important for the cytoplasmic localization and axonemal dynamics of intraflagellar transport proteins. Cilia 2, 13 (2013). 2046-2530-2-13 [pii] 10.1186/2046-2530-2-13

13 Gerondopoulos, A. et al. Planar Cell Polarity Eyector Proteins Inturned and Fuzzy Form a Rab23 GEF Complex. Curr Biol 29, 3323–3330.e3328 (2019). 10.1016/j.cub.2019.07.090

14 Gray, R. S. et al. The planar cell polarity eyector Fuz is essential for targeted membrane trayicking, ciliogenesis and mouse embryonic development. Nat Cell Biol 11, 1225–1232 (2009). ncb1966 [pii] 10.1038/ncb1966

15 Langousis, G. et al. Structure of the ciliogenesis-associated CPLANE complex. Sci Adv 8, eabn0832 (2022). 10.1126/sciadv.abn0832

16 Kim, S. K. et al. Planar Cell Polarity Acts Through Septins to Control Collective Cell Movement and Ciliogenesis. Science 329, 1337–1340 (2010). science.1191184 [pii] 10.1126/science.1191184

17 Zeng, H., Hoover, A. N. & Liu, A. PCP eyector gene Inturned is an important regulator of cilia formation and embryonic development in mammals. Dev Biol 339, 418–428 (2010). S0012-1606(10)00010-2 [pii] 10.1016/j.ydbio.2010.01.003

18 Heydeck, W. & Liu, A. PCP eyector proteins inturned and fuzzy play nonredundant roles in the patterning but not convergent extension of mammalian neural tube. Dev Dyn 240, 1938–1948 (2011). 10.1002/dvdy.22696

19 Cui, C., Chatterjee, B., Lozito, T., Zhang, Z. & Lo, C. W. Wdpcp, a PCP Protein Required for Ciliogenesis, Regulates Directional Cell Migration and Cell Polarity by Direct Modulation of the Actin Cytoskeleton. PLoS Biol. 11, e1001720 (2013).

20 Engelhardt, D., Marean, A., McKean, D., Petersen, J. & Niswander, L. RSG1 is required for cilia-dependent neural tube closure. Genesis 62, e23602 (2024). 10.1002/dvg.23602

21 Agbu, S. O., Liang, Y., Liu, A. & Anderson, K. V. The small GTPase RSG1 controls a final step in primary cilia initiation. J Cell Biol 217, 413–427 (2018). 10.1083/jcb.201604048

22 Toriyama, M. et al. The ciliopathy-associated CPLANE proteins direct basal body recruitment of intraflagellar transport machinery. Nat Genet 48, 648–656 (2016). 10.1038/ng.3558

23 Martín-Salazar, J. E. & Valverde, D. CPLANE Complex and Ciliopathies. Biomolecules 12 (2022). 10.3390/biom12060847

24 Bruel, A. L. et al. INTU-related oral-facial-digital syndrome type VI: A confirmatory report. Clin Genet 93, 1205–1209 (2018). 10.1111/cge.13238

25 Yakar, O. & Tatar, A. INTU-related oral-facial-digital syndrome XVII: Clinical spectrum of a rare disorder. Am J Med Genet A 188, 590–594 (2022). 10.1002/ajmg.a.62527

26 Zhang, W. et al. Expanding the genetic architecture and phenotypic spectrum in the skeletal ciliopathies. Hum Mutat 39, 152–166 (2018). 10.1002/humu.23362

27 Lopez, E. et al. C5orf42 is the major gene responsible for OFD syndrome type VI. Hum Genet 133, 367–377 (2014). 10.1007/s00439-013-1385-1

28 Shaheen, R. et al. Genomic analysis of Meckel-Gruber syndrome in Arabs reveals marked genetic heterogeneity and novel candidate genes. Eur J Hum Genet 21, 762–768 (2013). ejhg2012254 [pii] 10.1038/ejhg.2012.254

29 Saari, J., Lovell, M. A., Yu, H. C. & Bellus, G. A. Compound heterozygosity for a frame shift mutation and a likely pathogenic sequence variant in the planar cell polarity— ciliogenesis gene WDPCP in a girl with polysyndactyly, coarctation of the aorta, and tongue hamartomas. Am J Med Genet A 167a, 421–427 (2015). 10.1002/ajmg.a.36852

30 Steichen-Gersdorf, E., Gassner, I., Covi, B. & Fischer, H. Oral-facial-digital syndrome II. Transitional type between Mohr and Majewski syndrome: report of a new case with congenital stenosis of the trachea. Clin Dysmorphol 3, 245–250 (1994).

31 Zheng, D. et al. Predicting the translation eyiciency of messenger RNA in mammalian cells. bioRxiv, 2024.2008.2011.607362 (2024). 10.1101/2024.08.11.607362

32 Tabler, J. M. et al. Cilia-mediated Hedgehog signaling controls form and function in the mammalian larynx. Elife 6 (2017). 10.7554/eLife.19153

33 Abramson, J. et al. Accurate structure prediction of biomolecular interactions with AlphaFold 3. Nature 630, 493–500 (2024). 10.1038/s41586-024-07487-w

34 Walentek, P. & Quigley, I. K. What we can learn from a tadpole about ciliopathies and airway diseases: Using systems biology in Xenopus to study cilia and mucociliary epithelia. Genesis 55 (2017). 10.1002/dvg.23001

35 Rao, V. G. & Kulkarni, S. S. Xenopus to the rescue: A model to validate and characterize candidate ciliopathy genes. Genesis 59, e23414 (2021). 10.1002/dvg.23414

36 Srour, M. et al. Mutations in C5ORF42 cause Joubert syndrome in the French Canadian population. Am J Hum Genet 90, 693–700 (2012). S0002-9297(12)00096-1 [pii] 10.1016/j.ajhg.2012.02.011

37 Gerondopoulos, A., Langemeyer, L., Liang, J. R., Linford, A. & Barr, F. A. BLOC-3 mutated in Hermansky-Pudlak syndrome is a Rab32/38 guanine nucleotide exchange factor. Curr Biol 22, 2135–2139 (2012). 10.1016/j.cub.2012.09.020

38 Herrmann, E. et al. Structure of the metazoan Rab7 GEF complex Mon1-Ccz1-Bulli. Proc Natl Acad Sci U S A 120, e2301908120 (2023). 10.1073/pnas.2301908120

39 Onnis, A. et al. The small GTPase Rab29 is a common regulator of immune synapse assembly and ciliogenesis. Cell Death DiQer 22, 1687–1699 (2015). 10.1038/cdd.2015.17

40 Sharma, R. et al. The CPLANE protein Fuzzy regulates ciliogenesis by suppressing actin polymerization at the base of the primary cilium via p190A RhoGAP. Development 151 (2024). 10.1242/dev.202322

41 Yasunaga, T. et al. The polarity protein Inturned links NPHP4 to Daam1 to control the subapical actin network in multiciliated cells. J Cell Biol 211, 963–973 (2015). 10.1083/jcb.201502043

42 Megaw, R. et al. Ciliary tip actin dynamics regulate photoreceptor outer segment integrity. Nat Commun 15, 4316 (2024). 10.1038/s41467-024-48639-w

43 Kohli, P. et al. The ciliary membrane-associated proteome reveals actin-binding proteins as key components of cilia. EMBO Rep 18, 1521–1535 (2017). 10.15252/embr.201643846

44 Schrauwen, I. et al. FAM92A Underlies Nonsyndromic Postaxial Polydactyly in Humans and an Abnormal Limb and Digit Skeletal Phenotype in Mice. J Bone Miner Res 34, 375–386 (2019). 10.1002/jbmr.3594

45 Li, F. Q. et al. BAR Domain-Containing FAM92 Proteins Interact with Chibby1 To Facilitate Ciliogenesis. Mol Cell Biol 36, 2668–2680 (2016). 10.1128/mcb.00160-16

46 Burke, M. C. et al. Chibby promotes ciliary vesicle formation and basal body docking during airway cell diyerentiation. J Cell Biol 207, 123–137 (2014). 10.1083/jcb.201406140

47 Wang, C. et al. Centrosomal protein Dzip1l binds Cby, promotes ciliary bud formation, and acts redundantly with Bromi to regulate ciliogenesis in the mouse. Development 145 (2018). 10.1242/dev.164236

48 Lapart, J. A. et al. Dzip1 and Fam92 form a ciliary transition zone complex with cell type specific roles in Drosophila. Elife 8 (2019). 10.7554/eLife.49307

49 Siller, S. S., Burke, M. C., Li, F. Q. & Takemaru, K. Chibby functions to preserve normal ciliary morphology through the regulation of intraflagellar transport in airway ciliated cells. Cell Cycle 14, 3163–3172 (2015). 10.1080/15384101.2015.1080396

50 van Breugel, M., Rosa, E. S. I. & Andreeva, A. Structural validation and assessment of AlphaFold2 predictions for centrosomal and centriolar proteins and their complexes. Commun Biol 5, 312 (2022). 10.1038/s42003-022-03269-0

51 Wang, L. et al. FAM92A1 is a BAR domain protein required for mitochondrial ultrastructure and function. J Cell Biol 218, 97–111 (2019). 10.1083/jcb.201806191

52 Evans, R. et al. Protein complex prediction with AlphaFold-Multimer. bioRxiv, 2021.2010.2004.463034 (2022). 10.1101/2021.10.04.463034

53 Pylypenko, O., Hammich, H., Yu, I. M. & Houdusse, A. Rab GTPases and their interacting protein partners: Structural insights into Rab functional diversity. Small GTPases 9, 22–48 (2018). 10.1080/21541248.2017.1336191

54 Ostermeier, C. & Brunger, A. T. Structural basis of Rab eyector specificity: crystal structure of the small G protein Rab3A complexed with the eyector domain of rabphilin-3A. Cell 96, 363–374 (1999). 10.1016/s0092-8674(00)80549-8

55 Wang, W. J. et al. CEP162 is an axoneme-recognition protein promoting ciliary transition zone assembly at the cilia base. Nat Cell Biol 15, 591–601 (2013). 10.1038/ncb2739

56 Sang, L. et al. Mapping the NPHP-JBTS-MKS protein network reveals ciliopathy disease genes and pathways. Cell 145, 513–528 (2011). S0092-8674(11)00477-6 [pii] 10.1016/j.cell.2011.04.019

57 Zhao, H., Khan, Z. & Westlake, C. J. Ciliogenesis membrane dynamics and organization. Semin Cell Dev Biol 133, 20–31 (2023). 10.1016/j.semcdb.2022.03.021

58 Serres, M. P. et al. MiniBAR/GARRE1 is a dual Rac and Rab eyector required for ciliogenesis. Dev Cell 58, 2477–2494.e2478 (2023). 10.1016/j.devcel.2023.09.010

59 Insinna, C. et al. Investigation of F-BAR domain PACSIN proteins uncovers membrane tubulation function in cilia assembly and transport. Nat Commun 10, 428 (2019). 10.1038/s41467-018-08192-9

60 Sreekumar, V. & Norris, D. P. Cilia and development. Curr Opin Genet Dev 56, 15–21 (2019). 10.1016/j.gde.2019.05.002

61 Bruel, A. L. et al. Fifteen years of research on oral-facial-digital syndromes: from 1 to 16 causal genes. J Med Genet 54, 371–380 (2017). 10.1136/jmedgenet-2016-10443662

62. Sobreira, N., Schiettecatte, F., Valle, D. & Hamosh, A. GeneMatcher: a matching tool for connecting investigators with an interest in the same gene. Hum Mutat 36, 928–930 (2015). 10.1002/humu.22844

63 Richards, S. et al. Standards and guidelines for the interpretation of sequence variants: a joint consensus recommendation of the American College of Medical Genetics and Genomics and the Association for Molecular Pathology. Genet Med 17, 405–424 (2015). 10.1038/gim.2015.30

64 Torres, J. Z., Miller, J. J. & Jackson, P. K. High-throughput generation of tagged stable cell lines for proteomic analysis. Proteomics 9, 2888–2891 (2009). 10.1002/pmic.200800873

65 Shevchenko, A., Tomas, H., Havlis, J., Olsen, J. V. & Mann, M. In-gel digestion for mass spectrometric characterization of proteins and proteomes. Nat Protoc 1, 2856–2860 (2006). 10.1038/nprot.2006.468

66 Zybailov, B. et al. Statistical analysis of membrane proteome expression changes in Saccharomyces cerevisiae. J Proteome Res 5, 2339–2347 (2006). 10.1021/pr060161n

67 Ding, S. et al. Comparative Proteomics Reveals Strain-Specific β-TrCP Degradation via Rotavirus NSP1 Hijacking a Host Cullin-3-Rbx1 Complex. PLoS Pathog 12, e1005929 (2016). 10.1371/journal.ppat.1005929

68 Wood, C. W. et al. BAlaS: fast, interactive and accessible computational alanine-scanning using BudeAlaScan. Bioinformatics 36, 2917–2919 (2020). 10.1093/bioinformatics/btaa026

