## Supplementary material for "The human ciliopathy protein RSG1 links the CPLANE complex to transition zone architecture": Supp. Table 2

**Table 2. Statistics and N#s**

| **Figure 2. 2Way ANOVA of patient allele basal body localization** | | | | | | |
| --- | --- | --- | --- | --- | --- | --- |
| N# = | Embryos | Cells | BBs |  |  |  |
| WT | 5 | 25 | 1747 |  |  |  |
| S72P | 6 | 26 | 1725 |  |  |  |
| G114E | 6 | 30 | 2095 |  |  |  |
| D184W | 10 | 48 | 3302 |  |  |  |
| Tukey's multiple comparisons test | Mean Diff. | 95.00% CI of diff. | Below threshold? | Summary | Adjusted P Value |  |
| WT vs. S72P | 0.3659 | 0.1859 to 0.5459 | Yes | *** | 0.0002 |  |
| WT vs. G114E | 0.4044 | 0.2244 to 0.5844 | Yes | **** | <0.0001 |  |
| WT vs. D184W | 0.006654 | -0.1734 to 0.1867 | No | ns | 0.9995 |  |
| S72P vs. G114E | 0.03855 | -0.1305 to 0.2076 | No | ns | 0.9093 |  |
| S72P vs. D184W | -0.3592 | -0.5283 to -0.1901 | Yes | *** | 0.0001 |  |
| G114E vs. D184W | -0.3978 | -0.5669 to -0.2287 | Yes | **** | <0.0001 |  |
| Test details | Mean 1 | Mean 2 | Mean Diff. | SE of diff. | q | DF |
| WT vs. S72P | 0.6091 | 0.2432 | 0.3659 | 0.06193 | 8.355 | 14.00 |
| WT vs. G114E | 0.6091 | 0.2047 | 0.4044 | 0.06193 | 9.235 | 14.00 |
| WT vs. D184W | 0.6091 | 0.6024 | 0.006654 | 0.06193 | 0.1519 | 14.00 |
| S72P vs. G114E | 0.2432 | 0.2047 | 0.03855 | 0.05817 | 0.9372 | 14.00 |
| S72P vs. G114E | 0.2432 | 0.2047 | 0.03855 | 0.05817 | 8.732 | 14.00 |
| S72P vs. D184W | 0.2432 | 0.6024 | -0.3592 | 0.05817 | 9.670 | 14.00 |
| G114E vs. D184W | 0.2047 | 0.6024 | -0.3978 | 0.05817 | 8.355 | 14.00 |
| **Figure 3. 2Way ANOVA of IFT43 Rescue** | | | | | | |
| N# = | Embryos | Cells | BBs |  |  |  |
| Control | 6 | 22 | 3058 |  |  |  |
| Rsg1 KD | 6 | 29 | 1200 |  |  |  |
| +WT Rsg1 | 6 | 26 | 1584 |  |  |  |
| +S72P | 7 | 27 | 1072 |  |  |  |
| +G114E | 6 | 21 | 1113 |  |  |  |
| +D184W | 6 | 22 | 1257 |  |  |  |
| Tukey's multiple comparisons test | Mean Diff. | 95.00% CI of diff. | Below threshold? | Summary | Adjusted P Value |  |
| Rsg1 KD vs. WT Rsg1 | -0.3175 | -0.3452 to -0.2897 | Yes | **** | <0.0001 |  |
| Rsg1 KD vs. S72P | -0.01443 | -0.04415 to 0.01529 | No | ns | 0.7368 |  |
| Rsg1 KD vs. G114E | -0.009733 | -0.03911 to 0.01965 | No | ns | 0.9349 |  |
| Rsg1 KD vs. D184W | -0.05662 | -0.08509 to -0.02815 | Yes | **** | <0.0001 |  |
| WT Rsg1 vs. S72P | 0.3030 | 0.2742 to 0.3318 | Yes | **** | <0.0001 |  |
| WT Rsg1 vs. G114E | 0.3077 | 0.2793 to 0.3362 | Yes | **** | <0.0001 |  |
| WT Rsg1 vs. D184W | 0.2608 | 0.2335 to 0.2881 | Yes | **** | <0.0001 |  |
| S72P vs. G114E | 0.004697 | -0.02543 to 0.03482 | No | ns | 0.9978 |  |
| S72P vs. D184W | -0.04219 | -0.07170 to -0.01267 | Yes | *** | 0.0007 |  |
| G114E vs. D184W | -0.04688 | -0.07605 to -0.01772 | Yes | **** | <0.0001 |  |
| Test details | Mean 1 | Mean 2 | Mean Diff. | SE of diff. | q | DF |
| Control vs. Rsg1 KD | 1.000 | 0.3666 | 0.6334 | 0.009734 | 92.03 | 6221 |
| Control vs. WT Rsg1 | 1.000 | 0.6840 | 0.3160 | 0.008759 | 51.01 | 6221 |
| Control vs. S72P | 1.000 | 0.3810 | 0.6190 | 0.01011 | 86.59 | 6221 |
| Control vs. G114E | 1.000 | 0.3763 | 0.6237 | 0.009985 | 88.34 | 6221 |
| Control vs. D184W | 1.000 | 0.4232 | 0.5768 | 0.009575 | 85.19 | 6221 |
| Rsg1 KD vs. WT Rsg1 | 0.3666 | 0.6840 | -0.3175 | 0.009734 | 46.12 | 6221 |
| Rsg1 KD vs. S72P | 0.3666 | 0.3810 | -0.01443 | 0.01043 | 1.957 | 6221 |
| Rsg1 KD vs. G114E | 0.3666 | 0.3763 | -0.009733 | 0.01031 | 1.336 | 6221 |
| Rsg1 KD vs. D184W | 0.3666 | 0.4232 | -0.05662 | 0.009987 | 8.017 | 6221 |
| WT Rsg1 vs. S72P | 0.6840 | 0.3810 | 0.3030 | 0.01011 | 42.39 | 6221 |
| WT Rsg1 vs. G114E | 0.6840 | 0.3763 | 0.3077 | 0.009985 | 43.58 | 6221 |
| WT Rsg1 vs. D184W | 0.6840 | 0.4232 | 0.2608 | 0.009575 | 38.52 | 6221 |
| S72P vs. G114E | 0.3810 | 0.3763 | 0.004697 | 0.01057 | 0.6286 | 6221 |
| S72P vs. D184W | 0.3810 | 0.4232 | -0.04219 | 0.01035 | 5.762 | 6221 |
| G114E vs. D184W | 0.3763 | 0.4232 | -0.04688 | 0.01023 | 6.480 | 6221 |
| **Figure 4. 2Way ANOVA of Docking Rescue** | | | | | | |
| N# = | Embryos | Cells | BBs |  |  |  |
| Control | 6 | 23 | 4385 |  |  |  |
| Rsg1 KD | 6 | 23 | 2900 |  |  |  |
| +WT Rsg1 | 6 | 24 | 2538 |  |  |  |
| +S72P | 7 | 23 | 3030 |  |  |  |
| +G114E | 6 | 18 | 2464 |  |  |  |
| +D184W | 6 | 22 | 2933 |  |  |  |
| Tukey's multiple comparisons test | Mean Diff. | 95.00% CI of diff. | Below threshold? | Summary | Adjusted P Value |  |
| Rsg1 KD vs. WT Rsg1 | -1.110 | -1.379 to -0.8410 | Yes | **** | <0.0001 |  |
| Rsg1 KD vs. S72P | -0.1552 | -0.4251 to 0.1148 | No | ns | 0.5692 |  |
| Rsg1 KD vs. G114E | -0.3572 | -0.6272 to -0.08726 | Yes | ** | 0.0024 |  |
| Rsg1 KD vs. D184W | -0.6097 | -0.8797 to -0.3398 | Yes | **** | <0.0001 |  |
| WT Rsg1 vs. S72P | 0.9548 | 0.6849 to 1.225 | Yes | **** | <0.0001 |  |
| WT Rsg1 vs. G114E | 0.7528 | 0.4828 to 1.023 | Yes | **** | <0.0001 |  |
| WT Rsg1 vs. D184W | 0.5003 | 0.2303 to 0.7702 | Yes | **** | <0.0001 |  |
| S72P vs. G114E | -0.2020 | -0.4724 to 0.06838 | No | ns | 0.2698 |  |
| S72P vs. D184W | -0.4545 | -0.7249 to -0.1841 | Yes | **** | <0.0001 |  |
| G114E vs. D184W | -0.2525 | -0.5229 to 0.01788 | No | ns | 0.0828 |  |
| Test details | Mean 1 | Mean 2 | Mean Diff. | SE of diff. | q | DF |
| Control vs. Rsg1 KD | -0.4353 | -2.235 | 1.800 | 0.09404 | 27.07 | 492.0 |
| Control vs. WT Rsg1 | -0.4353 | -1.125 | 0.6900 | 0.09404 | 10.38 | 492.0 |
| Control vs. S72P | -0.4353 | -2.080 | 1.645 | 0.09436 | 24.65 | 492.0 |
| Control vs. G114E | -0.4353 | -1.878 | 1.443 | 0.09436 | 21.62 | 492.0 |
| Control vs. D184W | -0.4353 | -1.626 | 1.190 | 0.09436 | 17.84 | 492.0 |
| Rsg1 KD vs. WT Rsg1 | -2.235 | -1.125 | -1.110 | 0.09404 | 16.69 | 492.0 |
| Rsg1 KD vs. S72P | -2.235 | -2.080 | -0.1552 | 0.09436 | 2.326 | 492.0 |
| Rsg1 KD vs. G114E | -2.235 | -1.878 | -0.3572 | 0.09436 | 5.354 | 492.0 |
| Rsg1 KD vs. D184W | -2.235 | -1.626 | -0.6097 | 0.09436 | 9.139 | 492.0 |
| WT Rsg1 vs. S72P | -1.125 | -2.080 | 0.9548 | 0.09436 | 14.31 | 492.0 |
| WT Rsg1 vs. G114E | -1.125 | -1.878 | 0.7528 | 0.09436 | 11.28 | 492.0 |
| WT Rsg1 vs. D184W | -1.125 | -1.626 | 0.5003 | 0.09436 | 7.498 | 492.0 |
| S72P vs. G114E | -2.080 | -1.878 | -0.2020 | 0.09451 | 3.023 | 492.0 |
| S72P vs. D184W | -2.080 | -1.626 | -0.4545 | 0.09451 | 6.801 | 492.0 |
| G114E vs. D184W | -1.878 | -1.626 | -0.2525 | 0.09451 | 3.779 | 492.0 |
| **Figure 7. 2Way ANOVA of Ciliation %** | | | | | | |
| N# = | Raw |  |  |  |  |  |
| Control | 280 |  |  |  |  |  |
| Rsg1 Clone #1 | 333 |  |  |  |  |  |
| Rsg1 Clone #2 | 410 |  |  |  |  |  |
| Tukey's multiple comparisons test | Mean Diff. | 95.00% CI of diff. | Below threshold? | Summary | Adjusted P Value |  |
| Control vs. #11 | 67.19 | 58.83 to 75.55 | Yes | **** | <0.0001 |  |
| Control vs. #12 | 71.17 | 62.81 to 79.53 | Yes | **** | <0.0001 |  |
| #11 vs. #12 | 3.980 | -4.381 to 12.34 | No | ns | 0.3123 |  |
| Test details | Mean 1 | Mean 2 | Mean Diff. | SE of diff. | q | DF |
| Control vs. #11 | 81.04 | 13.85 | 67.19 | 2.346 | 40.50 | 4.000 |
| Control vs. #12 | 81.04 | 9.870 | 71.17 | 2.346 | 42.90 | 4.000 |
| #11 vs. #12 | 13.85 | 9.870 | 3.980 | 2.346 | 2.399 | 4.000 |
| **Figure 7. 2Way ANOVA Fam92a1 TZ intensity** | | | | | | |
| N# = | Raw |  |  |  |  |  |
| Control | 80 |  |  |  |  |  |
| Rsg1 Clone #1 | 80 |  |  |  |  |  |
| Rsg1 Clone #2 | 80 |  |  |  |  |  |
| Tukey's multiple comparisons test | Mean Diff. | 95.00% CI of diff. | Below threshold? | Summary | Adjusted P Value |  |
| Control vs. #11 | 0.5300 | 0.4328 to 0.6272 | Yes | **** | <0.0001 |  |
| Control vs. #12 | 0.5427 | 0.4455 to 0.6399 | Yes | **** | <0.0001 |  |
| #11 vs. #12 | 0.01275 | -0.08445 to 0.1100 | No | ns | 0.9483 |  |
| Test details | Mean 1 | Mean 2 | Mean Diff. | SE of diff. | q | DF |
| Control vs. #11 | 1.000 | 0.4700 | 0.5300 | 0.04108 | 18.24 | 158.0 |
| Control vs. #12 | 1.000 | 0.4573 | 0.5427 | 0.04108 | 18.68 | 158.0 |
| #11 vs. #12 | 0.4700 | 0.4573 | 0.01275 | 0.04108 | 0.4390 | 158.0 |
| **Figure 8. 2Way ANOVA Nphp1 TZ intensity** | | | | | | |
| N# = | Raw |  |  |  |  |  |
| Control | 80 |  |  |  |  |  |
| Rsg1 Clone #1 | 80 |  |  |  |  |  |
| Rsg1 Clone #2 | 80 |  |  |  |  |  |
| Tukey's multiple comparisons test | Mean Diff. | 95.00% CI of diff. | Below threshold? | Summary | Adjusted P Value |  |
| Control vs. #11 | 0.4215 | 0.3156 to 0.5274 | Yes | **** | <0.0001 |  |
| Control vs. #12 | 0.4894 | 0.3835 to 0.5954 | Yes | **** | <0.0001 |  |
| #11 vs. #12 | 0.06793 | -0.03799 to 0.1739 | No | ns | 0.2855 |  |
| Test details | Mean 1 | Mean 2 | Mean Diff. | SE of diff. | q | DF |
| Control vs. #11 | 1.000 | 0.5785 | 0.4215 | 0.04477 | 13.31 | 158.0 |
| Control vs. #12 | 1.000 | 0.5106 | 0.4894 | 0.04477 | 15.46 | 158.0 |
| #11 vs. #12 | 0.5785 | 0.5106 | 0.06793 | 0.04477 | 2.146 | 158.0 |
| **Figure 8. 2Way ANOVA Nphp1 BB intensity** | | | | | | |
| N# = | Raw |  |  |  |  |  |
| Control | 126 |  |  |  |  |  |
| Intu KO | 66 |  |  |  |  |  |
| Fuz KO | 52 |  |  |  |  |  |
| Tukey's multiple comparisons test | Mean Diff. | 95.00% CI of diff. | Below threshold? | Summary | Adjusted P Value |  |
| Control vs. Intu KO | 0.4039 | 0.2017 to 0.6061 | Yes | **** | <0.0001 |  |
| Control vs. Fuz KO | 0.5890 | 0.3673 to 0.8107 | Yes | **** | <0.0001 |  |
| Intu KO vs. Fuz KO | 0.1851 | -0.03656 to 0.4068 | No | ns | 0.1211 |  |
| Test details | Mean 1 | Mean 2 | Mean Diff. | SE of diff. | q | DF |
| Control vs. Intu KO | 1.000 | 0.5961 | 0.4039 | 0.08517 | 6.706 | 116.0 |
| Control vs. Fuz KO | 1.000 | 0.4110 | 0.5890 | 0.09338 | 8.921 | 116.0 |
| Intu KO vs. Fuz KO | 0.5961 | 0.4110 | 0.1851 | 0.09338 | 2.804 | 116.0 |
| **2Way ANOVA of docking rescue experiment (Supp. Figure 5)** | | | | | | |
| N#= | Cells | BBs |  |  |  |  |
| Control | 12 | 2201 |  |  |  |  |
| Jbts17 KD | 16 | 2181 |  |  |  |  |
| +WT Jbts17 | 16 | 2352 |  |  |  |  |
| +R1569* | 12 | 1712 |  |  |  |  |
| Tukey's multiple comparisons test | Mean Diff. | 95.00% CI of diff. | Below threshold? | Summary | Adjusted P Value |  |
| Control vs. Jbts17 KD | 1.840 | 1.587 to 2.093 | Yes | **** | <0.0001 |  |
| Control vs. + WT | 0.3700 | 0.1175 to 0.6225 | Yes | ** | 0.0011 |  |
| Control vs. +R1569* | 1.690 | 1.437 to 1.943 | Yes | **** | <0.0001 |  |
| Jbts17 KD vs. + WT | -1.470 | -1.723 to -1.217 | Yes | **** | <0.0001 |  |
| Jbts17 KD vs. +R1569* | -0.1500 | -0.4025 to 0.1025 | No | ns | 0.4181 |  |
| + WT vs. +R1569* | 1.320 | 1.067 to 1.573 | Yes | **** | <0.0001 |  |
| Test details | Mean 1 | Mean 2 | Mean Diff. | SE of diff. | q | DF |
| Control vs. Jbts17 KD | -0.06000 | -1.900 | 1.840 | 0.09775 | 26.62 | 297.0 |
| Control vs. + WT | -0.06000 | -0.4300 | 0.3700 | 0.09775 | 5.353 | 297.0 |
| Control vs. +R1569* | -0.06000 | -1.750 | 1.690 | 0.09775 | 24.45 | 297.0 |
| Jbts17 KD vs. + WT | -1.900 | -0.4300 | -1.470 | 0.09775 | 21.27 | 297.0 |
| Jbts17 KD vs. +R1569* | -1.900 | -1.750 | -0.1500 | 0.09775 | 2.170 | 297.0 |
| + WT vs. +R1569* | -0.4300 | -1.750 | 1.320 | 0.09775 | 19.10 | 297.0 |
| **2Way ANOVA of (Supp. Figure 7)** | | | | | | |
| N# = | Raw |  |  |  |  |  |
| Control | 21 |  |  |  |  |  |
| Intu clone | 4 |  |  |  |  |  |
| Fuz clone | 17 |  |  |  |  |  |
| Tukey's multiple comparisons test | Mean Diff. | 95.00% CI of diff. | Below threshold? | Summary | Adjusted P Value |  |
| Control vs. Intu KO | 54.41 | 38.80 to 70.02 | Yes | **** | <0.0001 |  |
| Control vs. Fuz KO | 47.36 | 38.94 to 55.78 | Yes | **** | <0.0001 |  |
| Intu KO vs. Fuz KO | -7.048 | -22.66 to 8.562 | No | ns | 0.4980 |  |
| Test details | Mean 1 | Mean 2 | Mean Diff. | SE of diff. | q | DF |
| Control vs. Intu KO | 51.39 | -3.022 | 54.41 | 6.145 | 12.52 | 19.00 |
| Control vs. Fuz KO | 51.39 | 4.026 | 47.36 | 3.314 | 20.21 | 19.00 |
| Intu KO vs. Fuz KO | -3.022 | 4.026 | -7.048 | 6.145 | 1.622 | 19.00 |
